# Biophysical characterization of two commercially available preparations of the drug containing *Escherichia coli* L-Asparaginase 2

**DOI:** 10.1101/2020.11.11.379065

**Authors:** Talita Stelling de Araújo, Sandra M. N. Scapin, William de Andrade, Maira Fasciotti, Mariana T. Q. de Magalhães, Marcius S. Almeida, Luís Maurício T. R. Lima

## Abstract

The hydrolysis of asparagine and glutamine by L-asparaginase has been used to treat acute lymphoblastic leukemia for over four decades. Each L-asparaginase monomer has a long loop that closes over the active site upon substrate binding, acting as a lid. Here we present a comparative study two commercially available preparations of the drug containing Escherichia coli L-Asparaginase 2, performed by a comprehensive array of biophysical and biochemical approaches. We report the oligomeric landscape and conformational and dynamic plasticity of *E. coli* type 2 L-asparaginase (EcA2) present in two different formulations, and its relationship with L-aspartic acid, which is present in Aginasa, but not in Leuginase. EcA2 shows a composition of monomers and oligomers up to tetramers, which is mostly not altered in the presence of L-Asp. The N-terminal loop of Leuginase, which is part of the active site is flexibly disordered, but gets ordered as in Aginasa in the presence os L-Asp, while L-Glu only does so to a limited extent. Ion-mobility spectrometry–mass spectrometry reveals two conformers for the monomeric EcA2, one of which can selectively bind to L-Asp and L-Glu. Aginasa has higher resistance to *in vitro* proteolysis than Leuginase, and this is directly related to the presence of L-Asp.

## 1. Introduction

L-Asparaginase is an amidohydrolase found in a variety of microorganisms, plants, and animals. It catalyzes the hydrolysis of L-asparagine (L-Asn) to L-aspartate (L-Asp) and ammonia. The enzyme is widely used for the treatment of several hematopoietic malignancies and is a crucial element in the treatment of acute lymphoblastic leukemia (ALL) in children, with increasing use in adult chemotherapy [1]. The L-asparaginase mechanism of action consists of serum L-Asn depletion since, unlike normal cells, leukemic cells are incapable of synthesizing this amino acid, leading to protein synthesis deficiency and cell death [2]. Most isoforms of L-asparaginases exhibit an L-glutaminase secondary activity, which has been demonstrated to be essential for full anti-leukemic efficacy (3,4). Despite its clinical relevance, several important toxic side effects attributed to serum L-glutamine (L-Gln) depletion have been observed in patients receiving L-asparaginase therapy, including immunosuppression, hepatotoxicity, encephalopathy, pancreatitis, and coagulation dysfunction. Therefore, engineering or identifying enzymes that retain asparaginase and reduced glutaminase activity has been claimed to help decrease toxic effects [5], which requires a detailed understanding of the structural basis involved in substrate recognition by L-asparaginases.

The crystal structure of asparaginase was first reported in 1993 for enzymes from ErA [6] and EcA2 [7], both bound to L-Asp. The active form of all bacterial L-asparaginases is described as homotetramers of about 130 to 140 kDa, usually referred to as a dimer of dimers, with four identical non-cooperative active sites formed at the intimate dimer interface. Asparaginase topology is well-conserved among the microorganisms and consists of two domains characterized by an α/β fold [8]. The N-terminal domain contains the catalytic site that, when occupied by the substrate, modulates the closure of a 20-amino acid flexible loop (residues 10 to 35 on mature EcA2), which acts as a lid over the active site. The structural integrity of this element is essential for asparaginase activity since its proteolytical removal leads to enzyme inactivation [9]. Evidence obtained from crystal structures of *E. chrytsanthemi, E. corotovora*, and *Wolinella succinogenes* L-asparaginases indicates that the flexible loop may be involved in substrate selectivity. The binding mode of different known ligands of L-asparaginases, such as L-Asp, L-Glu, D-Asp, and succinic acid monoamide is well conserved, with the binding of L-Glu accompanied by disorder of the flexible loop (17,18,19).

Several asparaginase preparations are currently available for clinical use, including *E. coli* L-asparaginase II (EcA2), either native or conjugated with polyethylene glycol (PEG-Asparaginase), as well as an isoform extracted from *Erwinia chrytsanthemi* (ErA), available for patients with allergies to *E. coli* products. These products share the same therapeutic mechanism but differ considerably in their pharmacokinetics and immunological properties. EcA2 and ErA are both characterized by low *in vivo* stability due to the production of neutralizing antibodies and their protease susceptibility, leading to a rapid decline in L-asparaginase serum activity. Indeed, EcA2 and ErA have half-lives of just 30 and 16 hours, respectively, meaning that several administrations are required for asparagine depletion [13]. Large variations in plasma activity have been associated with pronounced differences in the extent and persistence of L-Asn depletion, preventing significant leukemic cell death [14].

Despite the high remission rate achieved with the use of this enzyme, L-asparaginase based therapies are characterized by a high individual variation in pharmacokinetic and pharmacodynamic profile. More strikingly, independent reports have demonstrated that equivalent preparations of L-asparaginase from different manufactures show substantial differences in enzyme half-lives, plasmatic activity and immunogenic profile despite having a similar sequence and in vitro activity (8,9,10,11). Differences in the formulation of such a large and complex biotherapeutic drug can strongly impact the final biological product and compromise quality, safety, and efficacy with potentially radical clinical consequences [19]. Since the macromolecular architecture plays a role in determining function and stability, a deep knowledge of the physicochemical landscape of similar biopharmaceuticals can provide anticipation of the product quality and clinical outcomes.

In this study, we investigated the oligomeric organization, stability, and dynamics of EcA2 and its response to selective ligand binding. We used two brands of biosimilar EcA2 products, which further allowed the interpretation of the findings at light of the currently available information concerning their enzymatic activity in vivo and immunogenic profile (10,11).

## 2. Material and Methods

### 2.1. Materials

The EcA2 samples were kindly donated by the Martagão Gesteira Childcare and Pediatrics Institute (IPPMG-UFRJ), Rio de Janeiro, Brazil. We used lot #F140328A for Aginasa and lot #2017010101 for Leuginase. The calculated molecular weight of mature asparaginase II from *E. coli* (Uniprot P00805; amino acids 23-348) was 34,593.94 Da for average mass and 34,572.56 Da for monoisotopic mass. The lyophilized powder containing EcA2 product was dissolved in water for injection (WFI) to 2,500 IU/mL and kept at 4 °C–8 °C prior to use in measurements. We used a conversion factor of about 255 IU/mg EcA2, based on the total potency declared in the product leaflet (https://www.accessdata.fda.gov/drugsatfda_docs/label/2013/101063s5169lbl.pdf). No further corrections were performed. All other reagents were of analytical grade.

### 2.2. Liquid chromatography - Electrospray ionization (ESI) - Mass spectrometry analysis of asparaginase products

The asparaginase samples were diluted to 200 μg/mL in 0.1% formic acid in water. Each chromatographic run contained a total protein amount of 4 μg (20 μL protein solution per injection). Chromatographic analysis was performed in a Discovery BIO Wide Pore C18 column (15 cm x 1 mm, 5 μm, Supelco, USA) in a Prominence chromatographic system (Shimadzu, Japan) at 60°C and 100 μL/min mobile phase flux.

The elution was performed with a binary gradient (A - 0.1% formic acid in water; B – 0.1% formic acid in acetonitrile) as follows:

- 0–1.5 min, 10% B
- 1.5–38.5 min, 10%–60% B
- Wash for 5 min with 95% B
- Re-equilibration with 10% B for 5 min

The detection was performed with a Diode Array (SPD-M20A Prominence, Shimadzu Co., Japan) scanning from 250 to 400 nm, followed in line by a mass spectrometer Impact (Bruker Daltonics, Germany) equipped with an ESI source, under the following conditions: gas nebulizer at 1.2 bar, drying gas flow at 8.0 L/min, capillary at 4.5 kV, source at 200°C. The spectra were acquired in positive mode in the range of m/z 250 to 3,500. The chromatogram and spectra were analyzed with DataAnalysis 4.2 (Bruker Daltonics, Germany) and mMass softwares [20]. All analyses were performed at the Center for Biomolecular Mass Spectrometry (CEMBIO/UFRJ), Rio de Janeiro, Brazil.

### 2.3. Limited Proteolysis Assay

L-asparaginase samples were diluted to a final concentration of 280 μM and incubated at 25°C with 0.9 μg/ml of Proteinase K (Calbiochem, Cat# 539480) in the absence or presence of a 10-fold excess of L-Asp or L-Glu (2.8 mM). A sample was collected after different times of incubation, and the reaction was ended by heating at 100°C for 15 minutes in Laemmli loading buffer. The digestion profile was visualized by 15% SDS-PAGE stained with colloidal Coomassie blue. After 60 minutes, the reactions were frozen in liquid nitrogen and stored at -80°C for further analysis using liquid chromatography, Electrospray ionization, and mass spectrometry at the Center for Biomolecular Mass Spectrometry (CEMBIO/UFRJ), Rio de Janeiro, Brazil. LC-MS samples were thawed and diluted to a final concentration of 1.5 μM in 0.1 % formic acid in water and immediately injected in the LC-ESI-MS system. Each sample was submitted to two LC runs with different detection methods, one performed without fragmentation in the range of 250–3,500 m/z to obtain the molecular mass of the final products after limited proteolysis. For fragmentation experiments, spectra were recorded in the range of 50–1500 m/z using the data-dependent acquisition mode (DDA/Auto-MS) with a three-precursor ion per fragmentation cycle.

### 2.4. Protein Enzymatic Digestion, MALDI-TOF-MS/MS Analysis, and Peptide de novo Sequencing

After the intact protein mass analysis by both MALDI-TOF/MS and ESI Q-TOF/MS systems, we performed enzymatic digestion for both proteins with trypsin. Briefly, 100 μg of Leuginase and Aginasa were independently incubated with a solution containing 20 mmol/L triethylammonium bicarbonate (TEAB), 0.1 mmol/L Tris(2-carboxyethyl) phosphine (TCEP), 40 mmol/L chloroacetamide at 45 °C for 2 hours. The proteins were digested overnight with trypsin (1:50) (Promega, USA) in 1% (w/v). The tryptic peptides were cleaned by using ZipTip TM C18 resin and submitted to MALDI-TOF/MS analysis using a saturated mixture of α-cyano-4-hydroxycinnamic acid (CHCA), spotted on a MALDI sample plate, and dried at room temperature. Peptide fragmentation was obtained by MALDI-TOF-MS/MS experiments and the resulting data were analyzed manually using mMass software (37).

### 2.5. Size Exclusion Chromatography

SEC analysis was performed using a TSKgel G3000SW_XL_ column (300 x 7.8 mm - Supelco) to determine the EcA2 hydrodynamic radius. We used samples of Aginasa, Leuginase, and Leuginase supplemented with L-Asp at 320, 150, 50, and 10 μM in 10 mM sodium phosphate at pH 7.0. Proteins were separated at a flow rate of 1 mL/min and monitored by OD_280 nm_. Fractions of each peak were collected and analyzed using 15% SDS-PAGE stained with colloidal Coomassie blue to confirm EcA2 elution. The following globular proteins were used to fit a standard curve of elution profile versus hydrodynamic radius: thyroglobulin 86 Å (Sigma-Aldrich, Cat #T9145) bovine serum albumin–dimer 45.6 Å and monomer 36.2 Å (Sigma-Aldrich, Cat #A7906), Ovalbumin 30.5 Å (Sigma-Aldrich, Cat #A5503), and carbonic anhydrase 21.4 Å (Sigma-Aldrich, Cat #C7025).

### 2.6. Direct infusion ESI-traveling wave – ion mobility spectrometry – mass spectrometry (ESI-TWIM-MS)

The asparaginase stocks were diluted to varying concentrations in different solutions (either 0.1% formic acid or a buffer comprised of 50 mM ammonium acetate at pH 7.0, 10% acetonitrile in the absence or the presence of L-amino acids in acid form), as indicated in the respective figure caption. Due to the lower signal, measurements performed in ammonium acetate were accumulated for up to 18 min at a flow rate of 20 μL/min.

ESI-TWIM-MS measurements were performed in a Traveling Wave Ion Mobility Mass Spectrometer (TWIM-MS, Synapt G1 HDMS, Waters, UK). The samples were directly infused at 20 μL/min, using a positive ESI with a capillary voltage of 4.0 kV and N_2_g used as mobility gas at 0.4 bar. Data acquisition was conducted over the range of m/z 500–3,000 for 5 minutes. The mass spectrometer calibration was performed with 0.1 % phosphoric acid (v/v) in acetonitrile: H_2_O (1:1). Other typical instrumental settings followed a previous study [21]. Data were analyzed according to [22] using DriftScope (Waters Corporation, UK). All measurements were performed at the Laboratory of Macromolecules (LAMAC) and Life Science Metrology Division (DIMAV) of of Brazil’s National Institute of Metrology Standardization and Industrial Quality (INMETRO), Rio de Janeiro, Brazil.

### 2.7. Protein Crystallography

Crystals were obtained by sitting-drop vapor diffusion using 2 μL protein solution with 2 μL precipitant solution equilibrated against 80 μL precipitant solution per well in a Corning 3225 multiwell plate. We used sparse matrix screening kits (Hampton Research, USA) and the recently prepared EcA2 formulations. The co-crystallization of Leuginase with L-Asp was performed with a protein sample supplemented with 800 μM L-Asp (final concentration) from a 20 mM stock solution of amino acid in water. The crystals were obtained directly from the kit and used for data collection. Small needles/rods crystals were observed from assays with Leuginase within 5 min of setting up the crystallization screen, both in the presence and absence of supplemented L-Asp.

The preliminary characterization of the crystals was performed at room temperature with crystals imbibed in Apiezon N and the x-ray diffraction pattern using CuKα radiation (|μS 2.0, Incoatec) collected in a Photon II detector mounted in a D8-Venture diffractometer (Bruker AXS, Germany). Measurements were carried out at the Laboratory for X-ray Diffraction – LDRX, Physics Institute, Federal Fluminense University (UFF), Rio de Janeiro, Brazil.

Several unsuccessful attempts were made to obtain crystals for both EcA2 formulations in the same crystallization condition. After characterizing the components of the formulations, we became aware that Leuginase contained mannitol and polyethylene glycol (PEG), which might have influenced the results from different crystallization conditions. The best diffraction quality crystals were obtained in:

#### Leuginase with or without L-Asp

200 mM magnesium acetate, 20% PEG 3350 (Hampton Research PEG Ion I solution 25, “PEG ion I25”)

#### Aginasa

200 mM ammonium acetate, 20 % PEG 3350 (Hampton Research PEG Ion I solution 30, “PEG ion I30”)

Crystals were harvested from drops using 10 μm nylon CryoLoop™ (Hampton Research, USA) and flash-frozen under an N_2_g stream at 100 K and a flow rate of 10 L/h (Cryostream 700, Oxford Cryogenics, UK). Data collection was performed in the MX2 beamline from the Brazilian Synchrotron Light Laboratory (LNLS) at Brazil’s National Center for Research in Energy and Materials (CNPEM), Campinas, Brazil, using 1.459 Å (8.4991 keV) radiation. Diffraction datasets were collected on Pillatus 6M detector (Dectris) with the following parameters:

#### Aginasa

PEG ion Screen I #30, collection with exposure of 1 second/0.1 degree/image, 300 degrees (3,000 images) in total; structure solved by molecular replacement using 1IHD.pdb as a search model.

#### Leuginase supplemented with L-Asp

PEG Ion Screen I #25, full X-ray diffraction dataset collection with an exposure of 3 second/0.1 degree/image, 270 degrees (2,700 images) in total; structure solved by molecular replacement using 1IHD.pdb as a search model.

#### Leuginase apo

PEG Ion Screen I #25, full X-ray diffraction dataset collection with an exposure of 0.5 second/0.1 degree/image, 180 degrees (1800 images) in total; structure solved by molecular replacement using 1IHD.pdb as a search model.

The images were indexed, processed, and integrated with XDS [23], and the molecular replacement was performed with 1IHD.pdb using the Python-based MX2-autoProc pipeline (developed by Dr. Andrey F. Z. Nascimento, CNPEM). The solved structures were further refined and rebuilt with C.O.O.T. [24], Refmac [25], and Phenix [26] using the mature EcA2 sequence (UNIPROT P00805). Water molecules were added using C.O.O.T. Structural validation of the models performed with CCP4 Suite v.6.0.2 showed that almost all dihedral angles are within the favored range. Figures were generated with PyMOL (*PyMOL* Molecular Graphics System, Version 2.0, Schrödinger LLC). Selected crystallographic parameters are found in Table S3. The resulting models were deposited under pdb codes 6UOH (Aginasa, co-crystallized with L-Asp), 6UOD (Leuginase – apo), and 6UOG (Leuginase co-crystallized with L-Asp). Data collection and refinement statistics are found in Table S3.

The estimative of the intermolecular distances between EcA2 catalytic site residues and the ligand L-Asp as well as Val27 intramolecular distances was performed with MolMol V1.0.7 (Koradi et al. 1996) using a maximal distance of 5Å. The list of distances obtained with this analysis is summarized in Table S5 and S6.

### 2.8. NMR spectroscopy

Nuclear Magnetic Resonance (NMR) measurements were performed at 25°C on a 400 MHz Bruker Avance III spectrometer equipped with a broadband inverse Z-axis gradient probe (Bruker Spin Corp., GmbH). ^1^H chemical shifts were referenced to 3-(trimethyl-silyl)-1-propanesulfonic acid, sodium salt (DSS). The ^13^C chemical shifts were indirectly referenced to DSS by using the absolute frequency ratios. All NMR spectra were processed and analyzed with Topspin 4.0.6 (Bruker Spin Corp., GmbH).

The NMR samples used to characterize the EcA2 formulation products were ultrafiltrated using Amicon (Millipore) with a 3 kDa cut-off to remove the protein from the solution. The flow-through, supplemented with 10% deuterium oxide, was used to acquire 1D ^1^H spectra with water suppression using excitation sculpting with gradients. The 2D [^1^H,^13^C]-HSQC spectra were collected using the whole formulation with 10% deuterium oxide. The spectra obtained were compared with the ones acquired for standard solutions of mannitol, L-Asp, and PEG. For the quantitative analysis, we acquired several 1D ^1^H spectra with serial dilutions ranging from 1,000 to 62.5 μM of L-Asp and 20 to 1.25 mg/mL of mannitol. The peak areas were integrated and used to generate a standard curve with the concentration of each compound.

NMR samples used for the amino acid titration analysis were prepared with 160 μM of EcA2 (Leuginase) in 10 mM sodium phosphate at pH 7.0, with the addition of the respective amino acid, for the acquisition of 1D ^1^H NMR spectra. The intensities observed for Y25 H^ε^ and H^δ^ signals, at 6.7 and 7.0 ppm, were normalized against a resonance at approximately 6.8 ppm since no intensity variation was found for this signal in the presence of the substrates.

The STD-NMR experiments were performed at 298 K on a Bruker Avance III spectrometer operating at ^1^H frequencies of 600 MHz equipped with a triple-resonance, Z-axis gradient 5 mm probe. The EcA2 used for these experiments were previously desalted in a 5-mL HiTrap Desalting (GE Healthcare, USA) and eluted in 10 mM sodium phosphate at pH 7.0. The samples were prepared with a 500-fold excess of ligand (L-Asp or L-Glu 20 mM) over protein, 40 μM. The on-resonance spectrum was obtained with the saturation of protein signals at 0.87 ppm and the off-resonance spectrum at 10 ppm, using the Bruker standard parameters (STDDIFFGP.3). A spin-lock filter of 25 ms was used to suppress protein signals. A saturation time build-up curve was obtained to evaluate if an accumulation of saturated ligand molecules was occurring. The saturation times used were 0.5, 1.0, 1.5, 2.0, 2.5, 3.0, 3.5, and 4.0 s with recycling delay of 4.0 s. STD amplification factors (A_STD_) were determined using the following equation [27]:

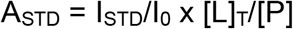

where [L]_T_/[P] is the molar ratio of the ligand in excess relative to the protein, and I_STD_/I_0_ is the ratio of the ligand signal integrals when the saturation is on-resonance (I_SDT_) or off-resonance (I_0_). The relative STD percentages were obtained using 2.5 s for saturation. The highest intensity of A_STD_ for each ligand was set to 100%, and all other STD signals were calculated relative to this.

### 2.9. Circular Dichroism

Circular dichroism experiments were performed on a Chirascan™ CD spectropolarimeter (Chirascan, Applied Photophysics, Leatherhead, UK) with a 100 μm path-length quartz cuvette. Far-UV spectra were recorded from 190 to 260 nm using EcA2 at 15 μM diluted in 10 mM sodium phosphate at pH 7.0 in the absence or presence of 1 mM L-Asp with a spectral resolution of 0.5 nm, a scan speed of 10 nm/min, and an average of three accumulations. The final data were obtained after subtraction of the buffer signal for background correction and are reported as molar residue ellipticity (deg.cm^2^.dmol^-1^), calculated as described in [28]. The thermal denaturation curve was monitored at 220 nm while the sample was heated from 25°C to 80°C in increments of 0.5°C using a rate of temperature change of 1°C/min. The data were processed using Prism 6 (GraphPad Software, USA). The melting temperature (T_m_), determined as the inflection point and slope of the sigmoidal curve, was calculated using a Boltzmann sigmoid equation.

### 2.10. Small-angle X-ray scattering (SAXS)

SAXS measurements were performed at 25°C in the D01A/SAS1 beamline from the LNLS/CNPEM equipped with a Pilatus 300 K detector following [29]. Samples were directly measured in a mica sample holder (1 mm optical path) and adequately subtracted for blank after total attenuation and photon counting. The samples were exposed for 300 sec at 1.488 Å, and no radiation damage was inferred when compared to shorter exposures. The scattering pattern was integrated with Fit2D [30], and ATSAS Suite 2.8.2 [31] was used in data analysis. The oligomeric distribution was calculated with *Oligomer* [32] using form factors calculated with *FFMaker* and the structural models designed from the EcA2 from Elspar (PDB ID 3ECA) using PyMOL in the q range of 0.022 A^-1^ to 0.48 A^-1^.

## 3. Results

### 3.1. The dissimilar formulations of EcA2 products

The analysis of the intact EcA2 products by LC-MS revealed a major peak in the chromatograms of Aginasa (Fig. 1A) and Leuginase (Fig. 1B), whose charge envelopes (Fig. 1C and 1D) allowed the estimation of their molecular mass at 34,590 Da and 34,576 Da, respectively (Fig. 1E). We have no objective explanation for the 15 Da difference between them. The leaflets of both products do not indicate any particular feature, such as post-translational or chemical modifications, which would be the most obvious explanations for such a difference.

**Figure 1.**
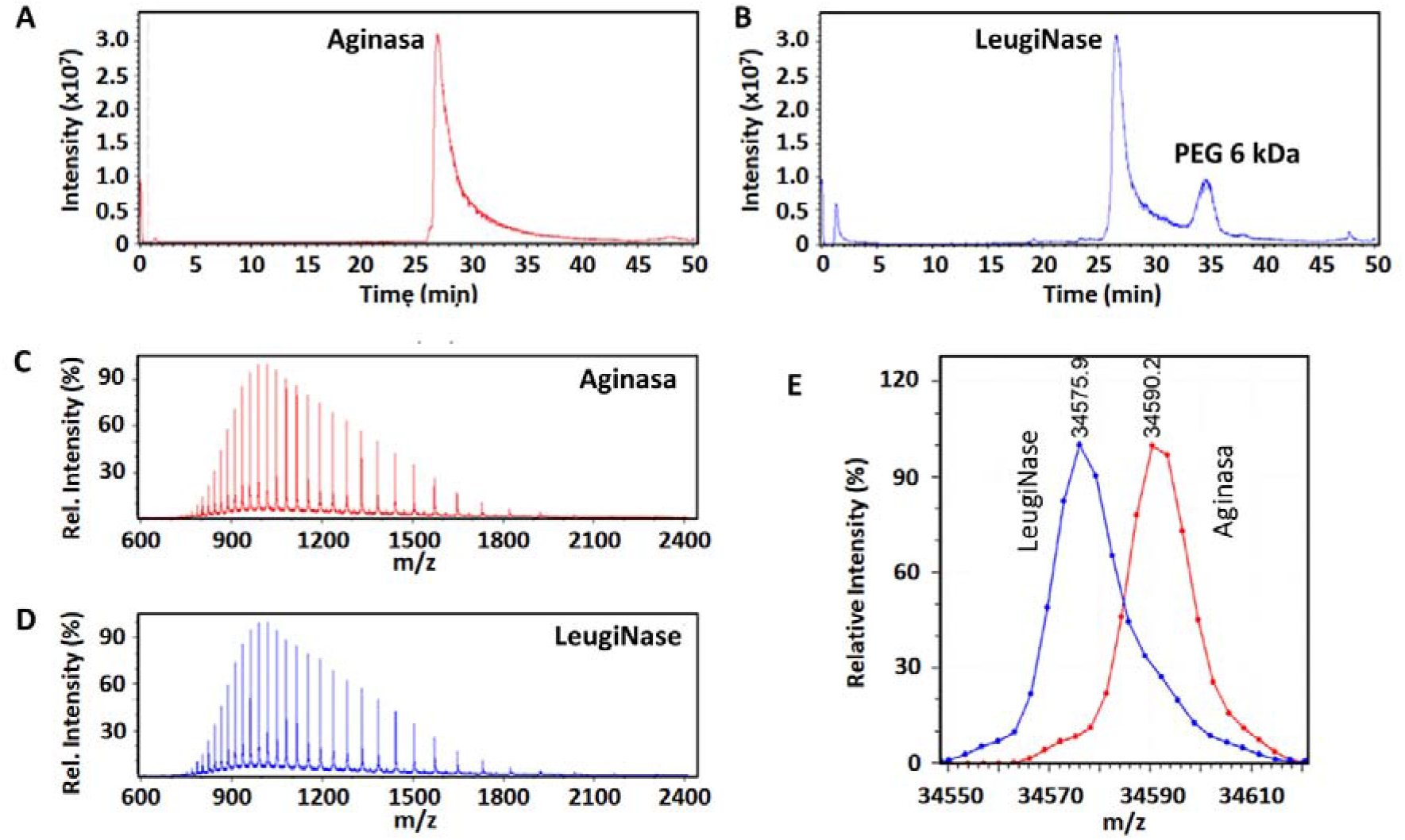
Characterization of asparaginase products by LC-ESI-MS. The EcA2 products Aginasa and Leuginase were evaluated by LC-ESI-MS. In the total ion current chromatograms, **A**. Aginasa and **B**. Leuginase showed a major peak corresponding to the respective intact proteins, and a peak corresponding to PEG with 6 kDa was found in Leuginase (Fig. S2). Charge envelopes **C**. and **D**. reveal the prevalence of monomers and minor contributions of dimers (smaller peaks with a thicker line between charged states belonging to monomers). **E**. The deconvoluted spectra revealed a mass of 34,590.2 Da for Aginasa and 34,575.9 Da for Leuginase.

In order to confirm the sequence identity of the protein we further performed enzymatic digestion of Aginasa and Leuginase with trypsin, followed by peptide identification by MALDI-TOF/TOF mass spectrometry. We found fragments corresponding to the predicted sequence of asparaginase for both Aginasa and Leuginase proteins (Fig. 1B), and the MS/MS spectra assignment for Aginasa and Leuginase [M+H]+ 2028.09 m/z allowed the corrected sequencing of the N-terminal 22 residues fragment (Fig. S1C).

We found a smaller peak in the Leuginase chromatogram (Fig. 1B), which, according to ESI-MS, corresponds to polyethylene glycol with an average mass of 6 kDa (Fig. S2). We performed a 2D [^1^H,^13^C]-NMR analysis and confirmed the presence of a PEG in Leuginase (Fig. 2B), as indicated by a major peak in its spectrum at about 3.6 ppm (^1^H) and 69.5 ppm (^13^C), which matches a test spectrum of PEG alone (Fig. 2D). In the NMR measurements, we identified other peaks at low field, which clearly correlate to mannitol (Fig. 2C). No peaks corresponding to PEG or mannitol were detected in Aginasa. However, other NMR peaks found in Aginasa but not in Leuginase correspond to free aspartate, at about 260 μM (ASP, which NMR could not distinguish between L and D forms) (Fig. 2A). The asparaginase product Elspar™,, which was used to obtain the first crystal structure of *E. coli* Asparaginase II (PDB ID 3ECA) [7], was reported to contain ASP, while its leaflet disclosed that its formulation included mannitol (https://www.accessdata.fda.gov/drugsatfda_docs/label/2002/aspamer080102LB.pdf). In opposite, Leuginase contains PEG and approximately 10 mM mannitol but no detectable ASP, as estimated from analytical curves (concentrations derived from quantitative NMR analysis shown in Fig. S3).

**Figure 2.**
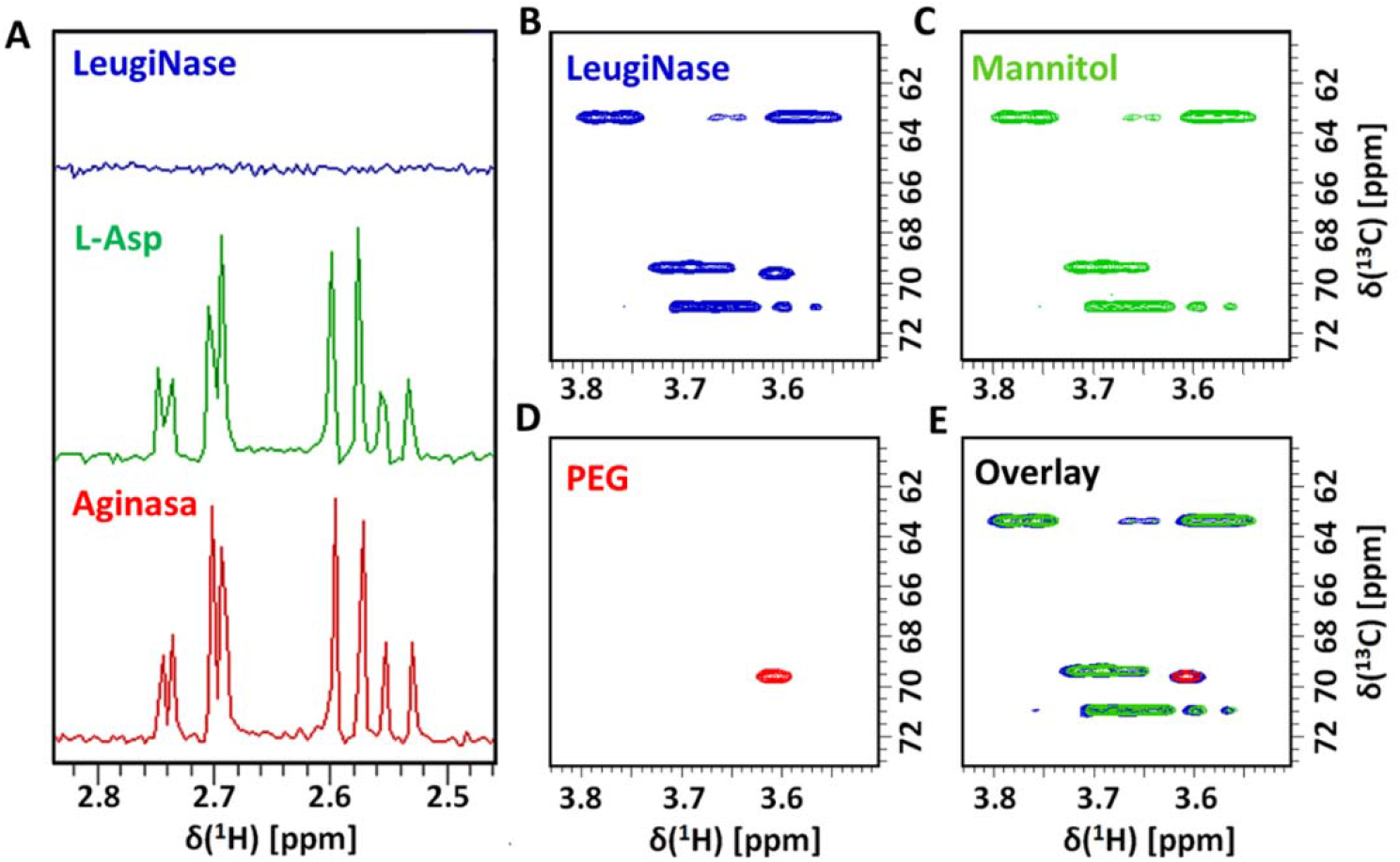
Identification of the constituents of EcA2 preparations using NMR **A**. 1D ^1^H NMR spectra of L-Asp (green) and ultrafiltrated EcA2 formulations, Aginasa (red) and Leuginase (blue). Spectra were collected in a 400 MHz spectrometer at 298 K. 2D [^1^H,^13^C]-HSQC spectrum of **B**. Leuginase total formulation, and standard solutions of **C**. mannitol and **D**. PEG. **E**. Overlay of the 2D spectra supports the presence of these constituents in the Leuginase formulation.

### 3.2. The thermodynamic stability of EcA2

To gain access to the overall conformation and stability of EcA2, we performed a thermal denaturation following changes in secondary structure by circular dichroism. We observed a large pre-transition plateau for both EcA2 products, with an abrupt transition accompanying a substantial change in the signal at 220 nm (Fig. 3). Both formulations presented similar Tm (67.52 ± 0.10°C for Aginasa and 67.87 ± 0.13°C for Leuginase). No apparent changes in their melting curves were observed when the experiment was performed in the presence of L-Asp (Tm of 66.78 ± 0.08°C for Aginasa plus L-Asp and 67.91 ± 0.06°C for Leuginase plus L-Asp). These thermodynamic data suggest a minor contribution of ligand binding and conformational change over broader folding energetics of the EcA2 against thermal denaturation. The most evident difference between the formulations was the slope for their thermal transition. Leuginase had slopes of 0.39 ± 0.02 and 0.44 ± 0.02, with and without the presence of 260 μM of L-Asp, respectively. These slopes were subtler for Aginasa (0.21 ± 0.02 and 0.22 ± 0.02, respectively).

**Figure 3.**
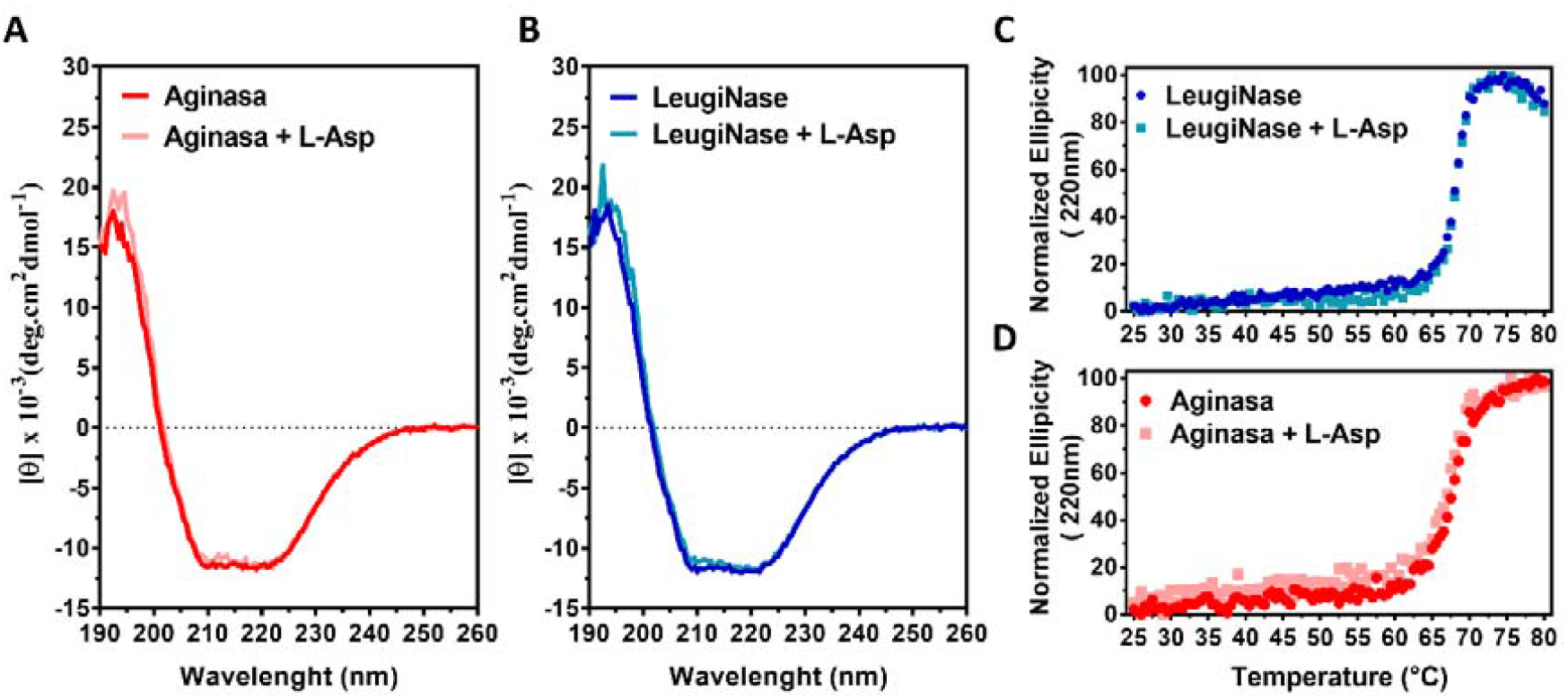
Thermal stability of EcA2. We monitored the thermal stability of EcA2 products, Aginasa and Leuginase, using circular dichroism (CD). CD spectra of **A**. Aginasa and **B**. Leuginase in the absence or presence of L-Asp showing only minor changes in absolute intensity. We tested the thermal stability of both **C**. Aginasa and **D**. Leuginase in the absence or presence of L-Asp. Sigmoidal fitting from 25°C to 80°C allowed the recovery of their transition midpoint and slope (shown in Table S1).

### 3.3. The oligomeric distribution of asparaginase by small-angle X-ray scattering (SAXS) and size-exclusion chromatography (SEC)

We performed SAXS measurements of the two EcA2 products to gain insight into the oligomeric distribution of EcA2. The scattering profile for each EcA2 product was similar between different vials reconstituted independently for Aginasa (Fig. 4A) and Leuginase (Fig. 4B). Attempts to deconvolute the oligomeric fraction from EcA2 Aginasa were not successful, even when oligomeric models of octamers or higher-order oligomers were included (Fig. S5). However, the deconvoluted scattering curves of Leuginase, revealed 81% tetramers, 10% trimers, 9% dimers, and no detectable monomers (Fig. 4E). Dilution of Leuginase up to 10 times showed no major effect on the presence of tetramers as a principal component in the oligomeric distribution of Leuginase (Fig. 4D).

**Figure 4.**
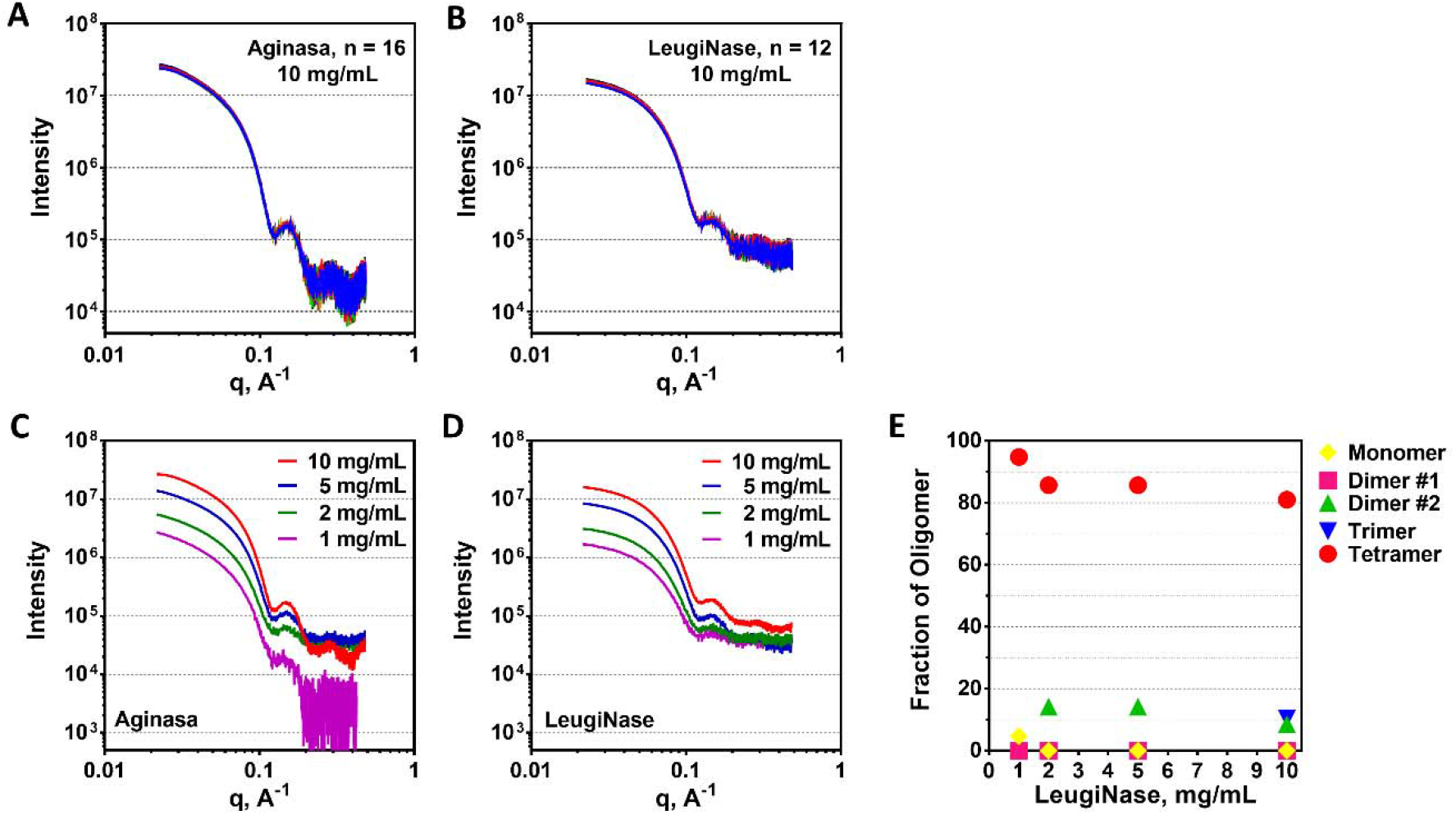
Small-angle X-ray scattering profile of EcA2 formulations. Each curve represents one sample at a different time from resuspension of the lyophilized product to data collection for **A**. Aginasa and **B**. Leuginase. Samples were stored at 4°C until measurements were performed at 25°C. Scattering profile of **C**. Aginasa and **D**. Leuginase at different protein concentrations. **E**. The fractions of monomer and the oligomers of Leuginase is shown for each concentration used.

To further characterize the oligomeric distribution of EcA2, we performed SEC using TSKgel G3000SW_XL_ (Fig. 5). We observed main peaks for Aginasa and Leuginase that were surrounded by minor peaks with distinct hydrodynamic radii, according to the standards used (Fig. **S**6 and Table S2). All marked peaks correspond to EcA2 for both products, as revealed by SDS-PAGE of the fraction collected from the chromatographed samples, suggesting that differences in the chromatographic peaks are likely to be related to oligomeric distribution or conformational fluctuations. The SEC profiles of both products are mostly not altered by dilution from 320 μM down to 10 μM or by the addition of L-Asp, confirming the SAXS findings for diluted samples (Fig. 4 and Fig. **S**7). The most noticeable difference is the relative abundance of fraction F3 and a deviation of the chromatogram of Aginasa. Given that the EcA2 substance is the major constituent of both products as indicated by LC-MS (Fig. 1), we hypothesize that the peak observed in Aginasa, with a retention time of 7.1 min, might be an indicator of higher-order assemblies. It is worth noting that the main peak of Leuginase presented a higher apparent elution time (8.7 min) in comparison to Aginasa (8.3 min).

**Figure 5.**
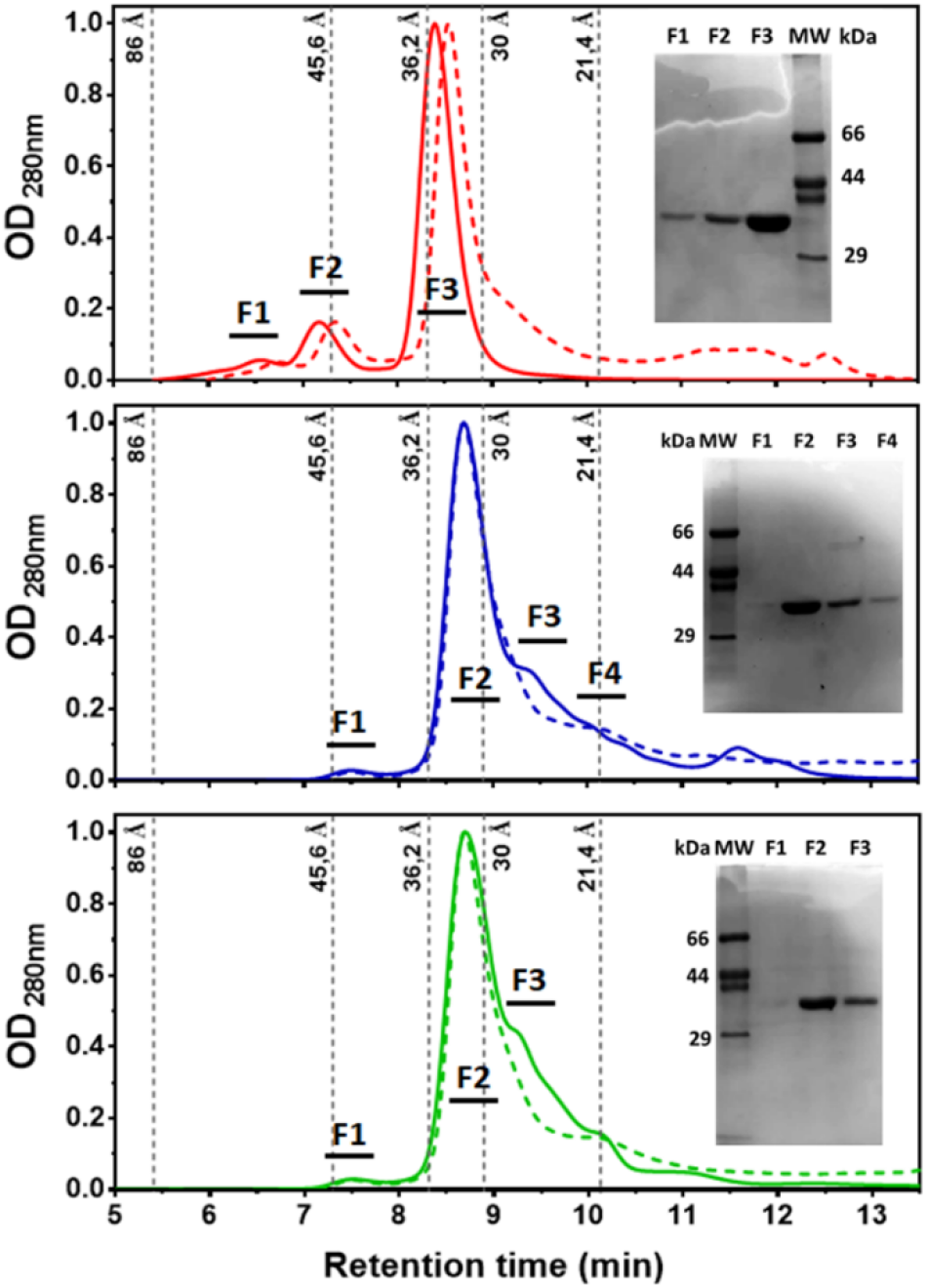
Size-exclusion chromatography analysis of Aginasa (red), Leuginase (blue), and Leuginase supplemented with 1 mM L-Asp (green). Analysis by SDS-PAGE 15% of the fractions obtained for each peak, confirming EcA2 elution. The hydrodynamic radius of standard globular proteins is indicated at their retention volume. Chromatograms with solid lines represent the sample at 320 μM, while dashed lines are for samples diluted to 10 μM. OD 280 nm was normalized to facilitate comparison of chromatographic runs.

### 3.4. The polymorphic distribution of Asparaginase and ligand interaction by ESI-TWIM-MS

We further explored the polymorphic distribution of EcA2 in solution using ESI-TWIM-MS. This method separates gas-phase ions based on their size, 3D shape, and ion-neutral interactions while the ions travel through a drift gas under the influence of an electric field. Studies have shown that this method is robust in determining the oligomeric and conformational distribution of proteins (20,21).

The deconvolution of the integrated spectra showed a mass difference of about 15 Da between the EcA2 products (Fig. 6), which independently confirms the results from intact LC-ESI-MS (Fig. 1). Both EcA2 products, Aginasa (Fig. 6A) and Leuginase (Fig. 6B), presented two populations in the ESI-TWIM-MS. The analysis of the stripped spectra corresponding to each of the two populations revealed their equivalence in molecular mass (Fig.s 6C and 6D). These data suggest the existence of EcA2 monomers in two defined populations of conformers.

**Figure 6.**
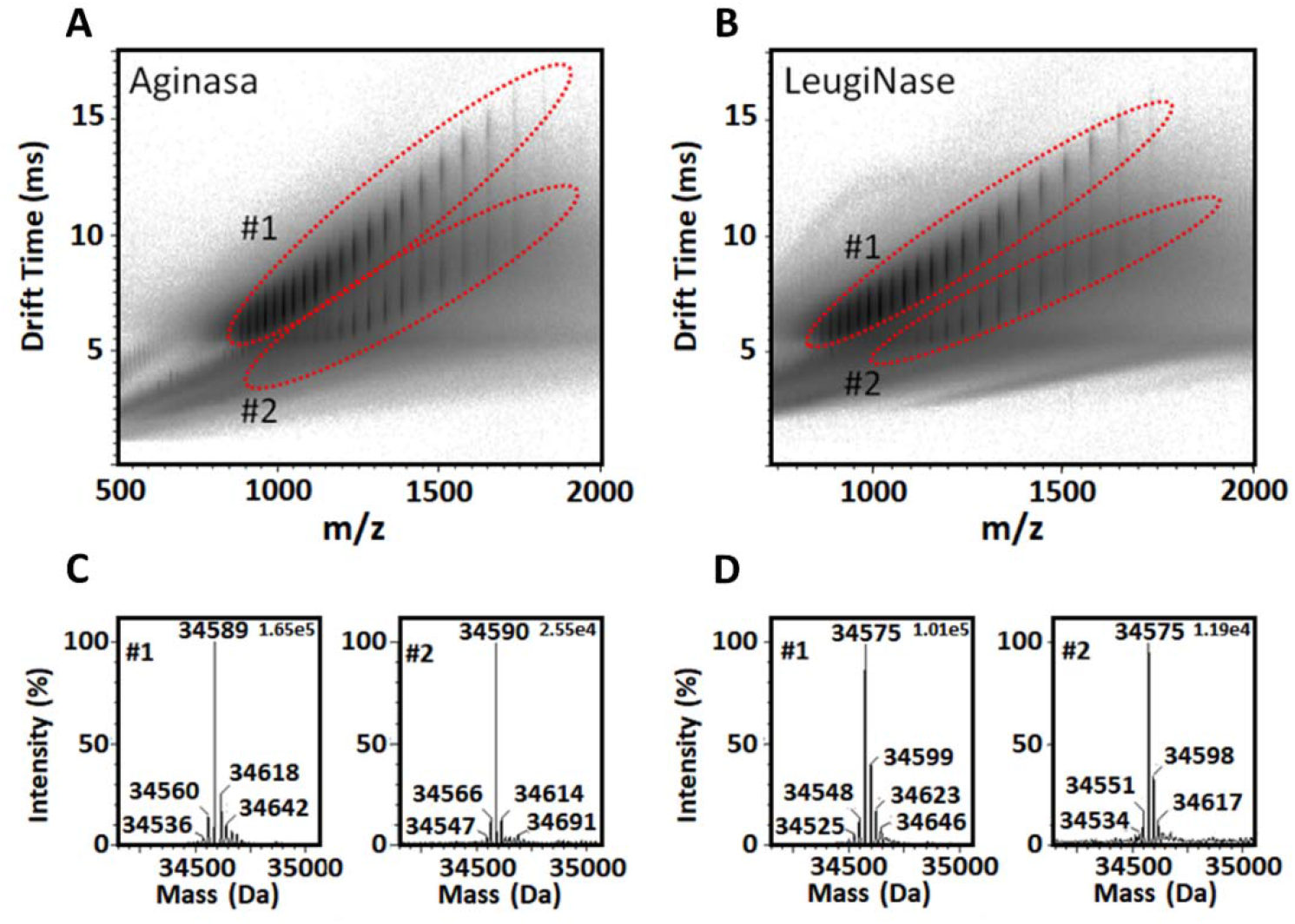
Multiple monomeric conformers of L-asparaginase as revealed by ion mobility spectrometry – mass spectrometry. **A**. Aginasa and **B**. Leuginase evaluated at 1 μM by ESI-IMS-MS in 0.1 % formic acid, showing two charge species envelopes (dotted red ellipses), corresponding to conformers with same molecular mass as shown by their deconvoluted spectra (**C** and **D**, respectively) using MaxEnt1 (MassLynx, Waters). Measurements performed in Synapt G1 (Waters) by direct injection.

ESI-TWIM-MS analysis of EcA2 in the presence of L-Asp or L-Glu revealed evidence for the amino acid-bound monomer (Fig. 7B and 7D) in varying charged states (M+15 to M+12) with a single distribution along the mobility axis (Fig. S8), indicating the distribution along with a well-defined population. Measurements performed in the presence of Gly showed a similar profile to the apo form (Fig. 7C), indicating the lack of detectable complex with this amino acid and ruling out non-specific EcA2: L-aa interaction by adduct effect. Collectively, these data demonstrate the selective nature of ligand interaction with EcA2 monomers.

**Figure 7.**
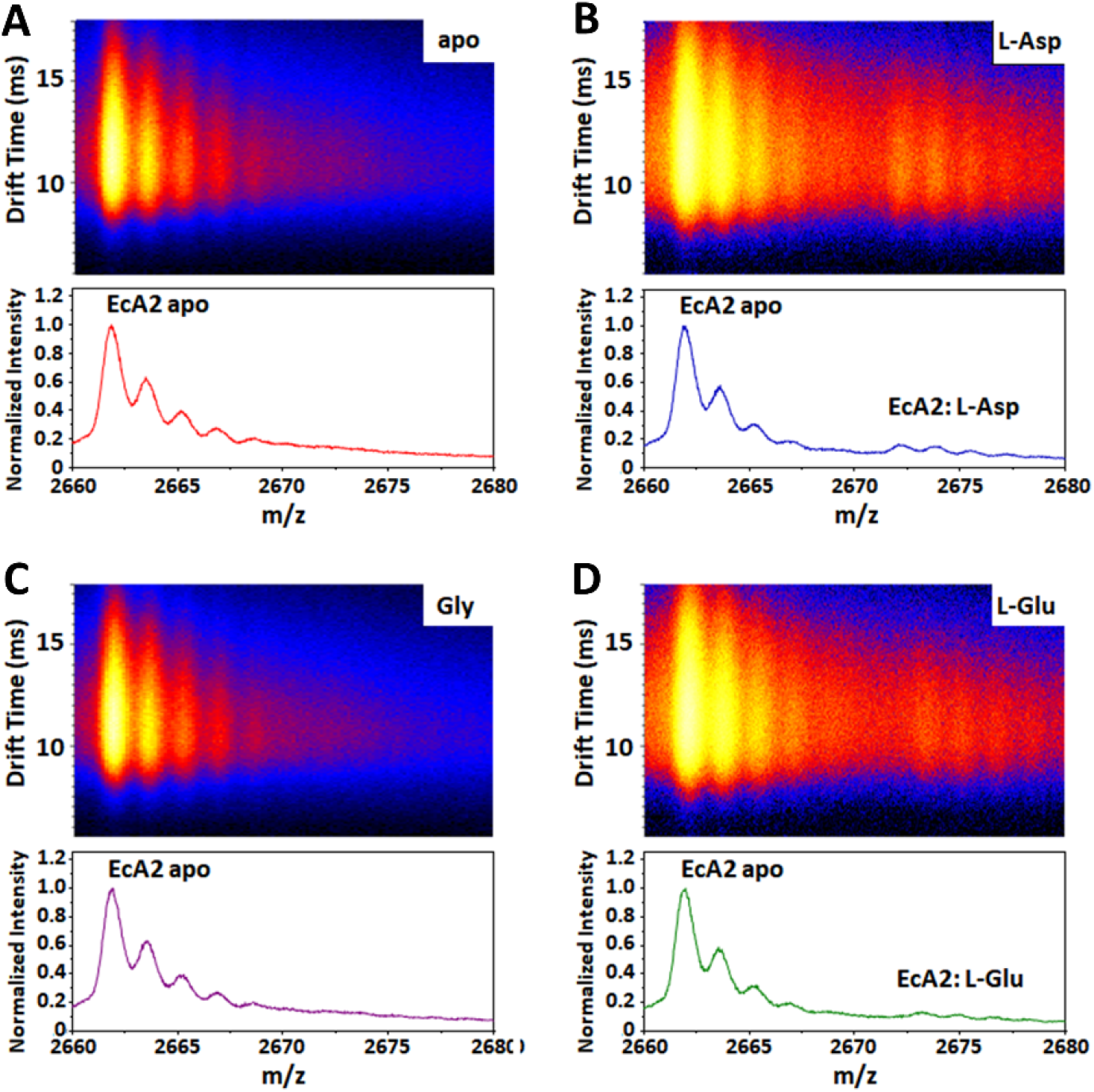
Mapping interaction of amino acid with EcA2 by mass spectrometry. The ESI-IMS-MS spectra of EcA2 were measured **A**. in the absence or the presence of **B**. L-aspartic acid, **C**. glycine, or **D**.L-glutamic acid. Only the M+13 charged state is shown for each sample, revealing the peaks corresponding to the free and the amino acid-bound forms of the protein and salt adducts (most likely Na). Other charged states are illustrated in Fig. S8, indicating similar results.

### 3.5. The structural basis for amino acid interaction with the EcA2 in the crystal phase

To gain more detailed insights on the structural basis of EcA2 interaction with ASP, we pursued the elucidation of the crystal structure of both EcA2. We solved the structures by obtaining diffracting quality crystals. The monomeric units within their asymmetric units were similar between the three structures, with overall RMSD of 1.2 Å or lower (Table S4). An electron density blob was observed within the catalytic site of Aginasa, which we attributed to L-Asp since NMR indicated the presence of this amino acid in the formulation (Fig. 2). In contrast, Leuginase was found in an ‘open’ conformation, without ligands in the active site and with only part of the N-terminal active site lid sufficiently visible for modeling, while Aginasa showed a well-ordered lid (Fig. 8). Attempts to crystallize Leuginase supplemented with L-Asp, resulted in cocrystallization of EcA2 with L-Asp in the active site and showing a closed lid, in close identity to the Asp-bound EcA2 from Aginasa (Fig. 8 and Table S4). These data demonstrate that EcA2 is structurally responsive to the interaction with L-Asp and the resulting consolidated crystal structural model of the two formulations are equivalent.

**Figure 8.**
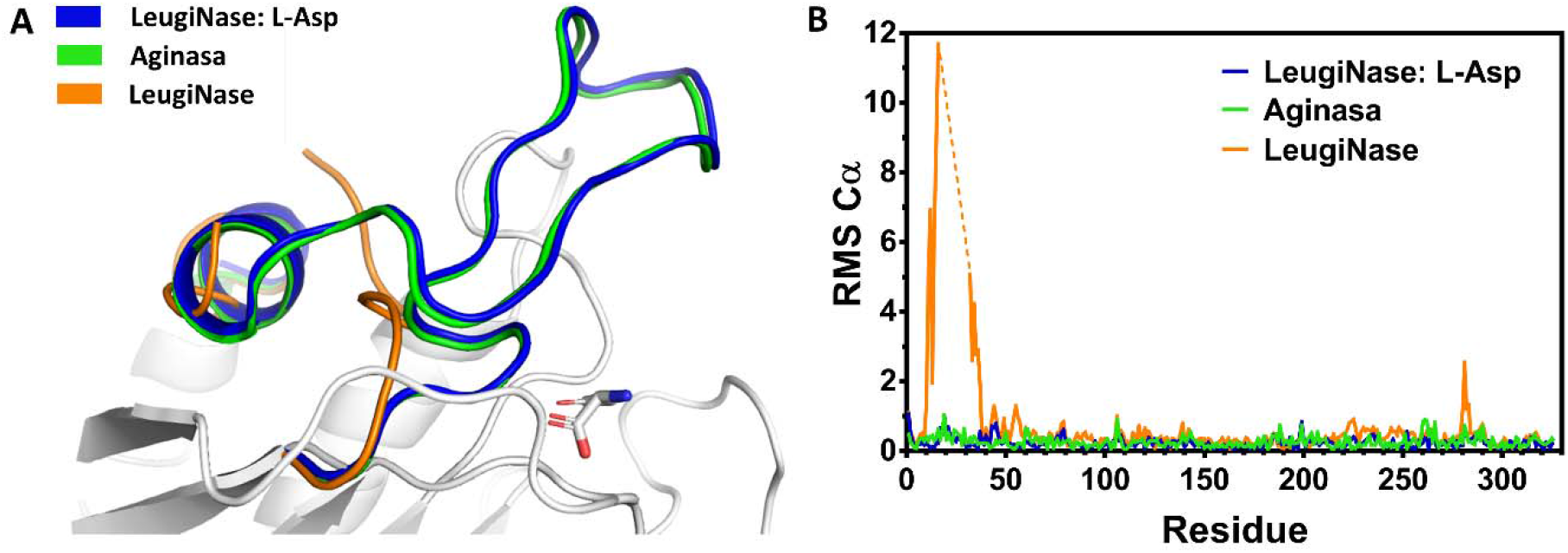
Crystallographic analysis of EcA2. Aginasa and Leuginase were subjected to crystallization and x-ray diffraction, and the structure was solved by molecular replacement. Aginasa (PDB Id.: 6UOH) showed two monomers in the asymmetric unit (A.U.) with bound L-Asp in the closed active site. In comparison, apo Leuginase (PDB Id.: 6UOD) showed four monomers in the A.U. and open active site with undefined lid. The co-crystallized Leuginase: L-Asp (PDB Id.: 6UOG) resulted in eight monomers in the A.U. with a closed lid over the amino acid-bound active site. **A**. Detailed view of the active-site lid, indicating the superposition between Asp-Aginasa and Asp-Leuginase. **B**. RMSD plot of the superposition of the crystal structures obtained, demonstrating the similarity in the spatial distribution along with the protein, except for the active-site lid (Residues G10 to N37) of the apo form of Leuginase. Data collection and refinement statistics can be found in the Table S3.

### 3.6. The dynamics of the active site lid using NMR

NMR revealed further insights on the dynamics of the active site lid upon binding to amino acids. 1D ^1^H-NMR spectra of the apo form of EcA2 (Leuginase) revealed two sharp peaks at 6.7 ppm and 7.0 ppm and three broad peaks at 7.7-8.4 ppm, not observed in the holo form of EcA2 (Aginasa) (Fig. 9A). The two sharp peaks were previously attributed to H^ε^ and H^δ^ of the aromatic ring of Tyr25, which belongs to the loop region comprising the lid over the active site of EcA2, and has been shown to be detectable in the apo form due to high conformational flexibility [35]. According to this interpretation, in case this loop gets more rigid, the peaks would get broader and fade out within the other broad signals in the spectra. Indeed, as observed in the EcA2 structure with bonded L-Asp, these signals disappear indicating the N-terminal loop closure in the presence of L-Asp or L-Asn. These changes were not observed when L-Glu or L-Gln were added to EcA2, indicating the selectivity of EcA2 for L-Asn and L-Asp for the induction of conformational changes in the loop region comprising the Tyr25, ultimately resulting in a closed form of the active site. The spectra do not present clear indication of alteration in the equilibrium of oligomerization of EcA2 in any of these samples, which would be represented by a global change in the line widths (Fig. **S**9). Instead, for example, there are peaks at 6.2, 6.6, 7.2 and 7.6 ppm, which mostly did not change their width.

**Figure 9.**
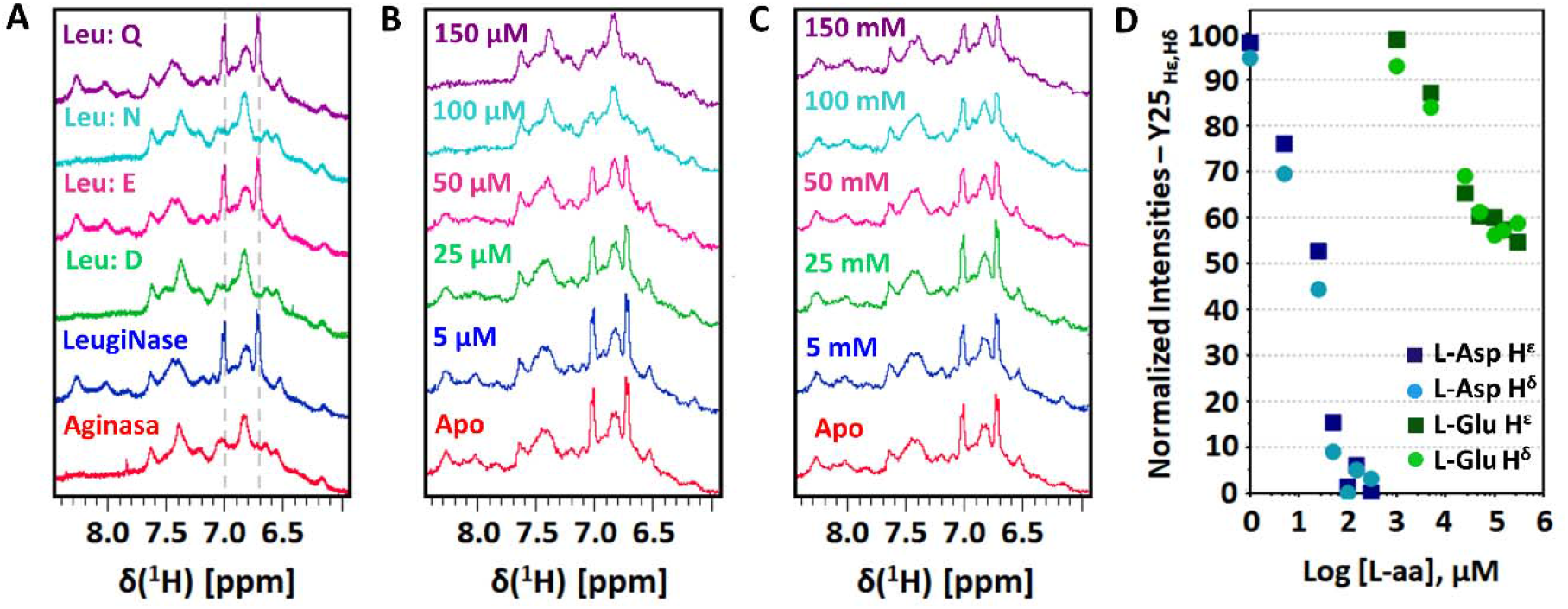
Monitoring Y25 dynamics upon L-aa binding by NMR. **A**. Multiple displays of 1D ^1^H NMR spectra of Aginasa and Leuginase whole formulation (320 μM) and Leuginase added with 1 mM of different L-amino acids indicated with the one-letter code above each spectrum. For each experiment, a total of 512 scans with 2 s of relaxation delay were collected in a 400 MHz spectrometer at 298 K. In the presence of L-Asp and L-Asn, the Tyr25 pair of sharp signals attributed to H^ε^ and H^δ^ (marked with a dotted gray line) disappear, indicating a loss of flexibility. Multiple displays of NMR-titration analysis of 160 μM Leuginase supplemented with increasing concentrations of **B**. L-Asp or **C**. L-Glu. Concentration of L-aa used for the acquisition of each spectrum is indicated in the figure. For each experiment, a total of 256 scans with 2 s of relaxation delay were collected in a 400 MHz spectrometer at 298K. **D**. The quantitative analysis of the NMR-titration.

To further investigate the selectivity of EcA2 for L-Asn/L-Asp over L-Gln/L-Glu and their ability to induce a conformational response, we performed titration analyses monitoring the aromatic resonances attributed to Tyr25 of EcA2 by 1D ^1^H-NMR. The titration with L-Asp (Fig. 9B) indicated that this ligand restricted the loop flexibility in an EcA2:L-Asp ratio of about 1:1, evidenced by the disappearance of a pair of sharp Tyr25 signals. When EcA2 was titrated with L-Glu (Fig. 9C), a significant change in Tyr25 resonances could only be detected with an EcA2:L-Glu ratio of over 1:300. Even with a 1:1800 proportion of ligand, the observable intensities of Tyr25 resonances were approximately 50% of the initial intensity prior to the addition of the ligand (Fig. 9D). This demonstrates the inability of L-Glu to induce conformational changes in EcA2 to the same extent as L-Asp.

### 3.7. L-Asp protects EcA2 against limited proteolysis of the active-site lid

To further evaluate the selective effect of amino acid-binding on the active site lid in the presence of ligand binding in the catalytic site, we tested the influence of L-Asp or L-Glu over the propensity of EcA2 for limited proteolysis. The resulting products from the proteolysis were subjected to identification by LC-ESI-MS, which revealed two major chromatographic peaks as proteolytic products, one corresponding to an ion equivalent in mass to the N-terminal residues 1-23 of EcA2 and the other to the remaining part of the protein (Fig. S8).

The EcA2 was incubated with proteinase-K in the absence and presence of L-Asp or L-Glu, and aliquots were taken at varying time intervals and subjected to SDS-PAGE. The EcA2 product Leuginase showed a time-dependent decrease in the main protein band corresponding to the unaltered protein with the progressive increase of a band with higher electrophoretic mobility (Fig. 10). The EcA2 product from Aginasa, which was found to contain an equimolar amount of ASP, showed slower proteolysis kinetics. Supplementation of both both Aginasa and Leuginase with further amount of L-Asp resulted in a decreased rate of degradation when compared to the assay conducted in the original EcA2 preparations. No protection was detected when the limited proteolysis was conducted in the presence of L-Glu. These data independently confirm the selective nature of ligand binding to the active site and its influence on the dynamics of the active site lid.

**Figure 10.**
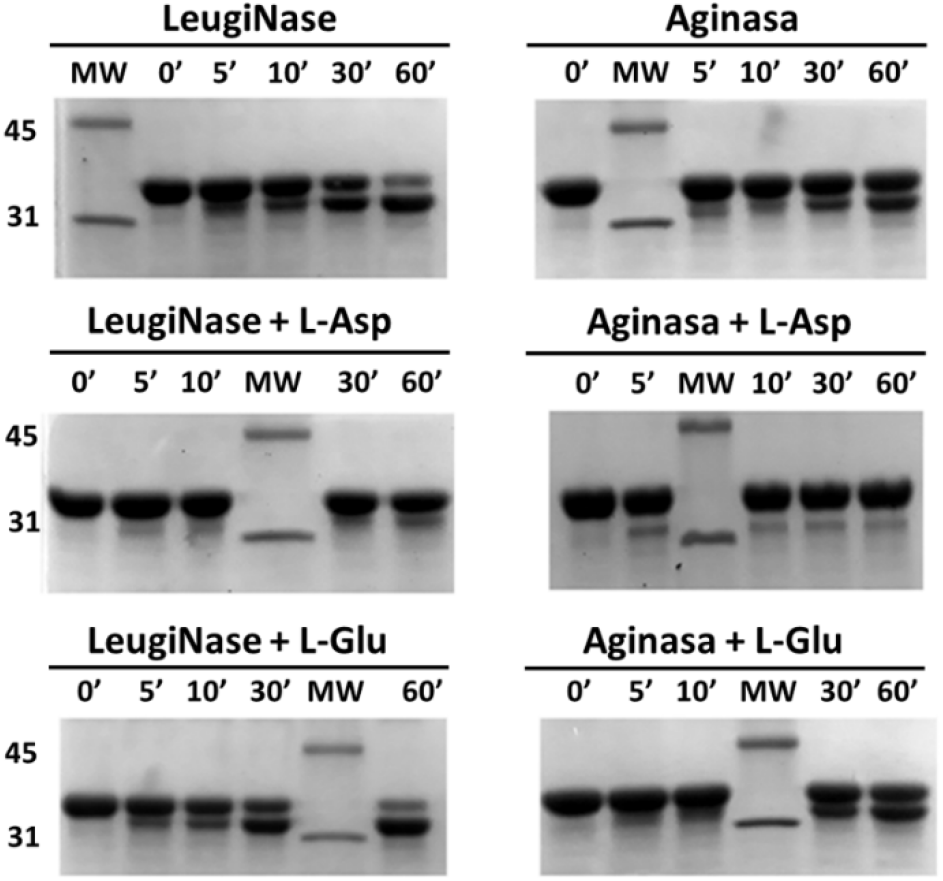
L-Asp protects EcA2 against limited proteolysis. SDS-PAGE 15% analysis of Aginasa and Leuginase (280 μM), with and without L-Asp or L-Glu addition (2.8 mM), after incubation with Proteinase K among indicated time. Two bands of the molecular weight (MW) ladder are indicated on the left (31 and 45 kDa).

### 3.8. Contact points and affinity of amino acid interaction with EcA2 in solution by STD-NMR

In addition to our evaluation of the selective nature of ligand binding on the dynamics of the active site lid of EcA2, we used STD-NMR to characterize these effects on the ligand. This method allows the recognition of the binding epitope on the ligand through the efficiency of saturation transfer from the protein that is saturated specifically by a long radio-frequency pulse. The efficiency of the saturation transfer is quantitatively expressed by the amplification factor (A_STD_). The intensity of A_STD_ characterizes the average number of transient contacts (≤ 5 Å) of the ligand per molecule of receptor in a given saturation time [27].

The plot of the A_STD_ at different saturation times obtained for L-Asp demonstrates that H^α^ exhibits a remarkably stronger A_STD_ than H^β2^ and H^β3^ (Fig. 11A). The relative STD percentages indicate that the H^α^ is closer to EcA2, while the H^β^ is more distant.

**Figure 11.**
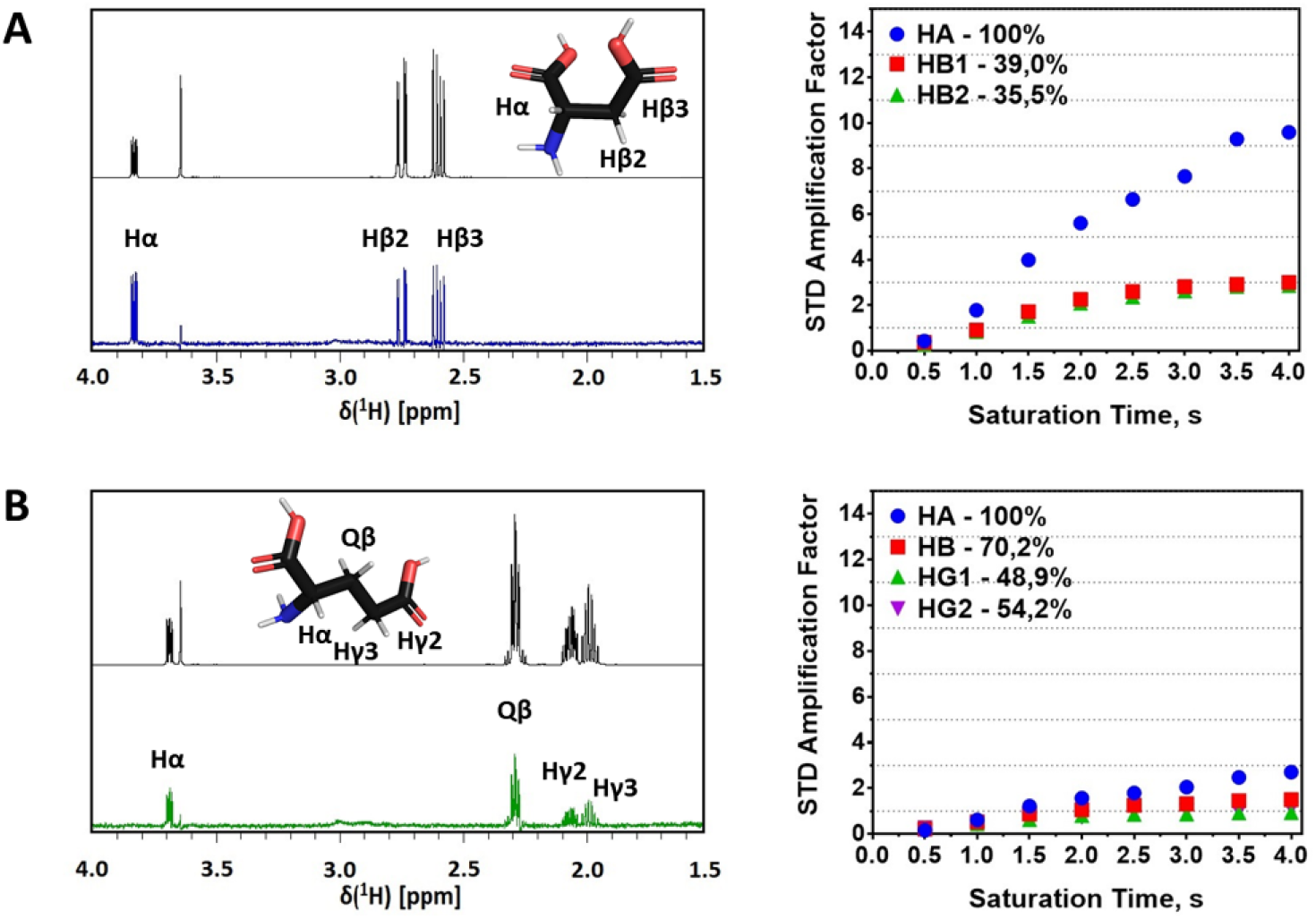
STD-NMR analysis reveal differences in substrate recognition of EcA2 for different ligands. The reference spectrum acquired for each ligand is in black, and the corresponding STD-NMR spectrum is colored in blue for **A**. L-Asp and green for **B**. L-Glu. The chemical structure of amino acids, BMRB entry bmse000031 (L-Asp) and bmse000037 (L-Glu), with assigned protons presented in each spectrum. The STD amplification factor as a function of saturation time is represented alongside each spectrum with relative STD percentages indicated for each assigned proton.

Additionally, we conducted STD-NMR experiments for L-Glu as an attempt to better describe the mechanism of EcA2 ligand recognition (Fig. 11B). Despite presenting a weaker STD signal due to a higher *K*_M_ for this ligand [36], the A_STD_ plot obtained for L-Glu shows relative STD percentages of 100%, 70.2%, 48.9%, and 54.2% for H^α^, H^β^, H^γ2^, and H^γ3^, respectively, indicating a weaker and more isotropic interaction of L-Glu in comparison to L-Asp.

## 4. Discussion

### 4.1. Structural and Dynamic Aspects

Asparaginase has long been described as a specific hydrolase for L-Asn, although the enzyme activity is also reported for L-Gln. In fact, our data reveal the propensity of EcA2 to bind to L-Asp and L-Glu, though not to Gly (Fig. 7, ESI-TWIM-MS). L-Asp induces conformational changes to close the active site lid (Fig. 8 and 9). Furthermore, L-Asp, but not L-Glu, is selective in changing the active site dynamics, as shown in our study by Tyr25 modulation (Fig. 9), limited proteolysis (Fig. 10), and binding measured by STD-NMR (Fig. 11). Specific ligand binding is thus a process that involves the final consolidated state, overall affinity, and kinetic processes.

EcA2 has been described as a tetramer (17,28). While a tetramer can be observed by SAXS (Fig. 4), as well as by crystallographic organization, our ESI-TWIM-MS data show the selective interaction of L-Asp and L-Glu with monomeric EcA2. Interestingly, the abundance of a tetramer is not apparent in the SEC-HPLC analysis (Fig. 5), where the main species has a calculated hydrodynamic radius of 31.2–34.8 Å (Table S2), which is most likely indicative of a dimeric state of EcA2. The interaction of EcA2 with ligands as monomers could be a result of enzymatic activity *in vivo* observed for hours after injection; this causes the extensive dilution of the protein and could favor its dissociation. However, the *K*_d_ of the tetramer and dimer in such biological milieus is unknown.

EcA2 has an N-terminal flexible loop that can form a species of lid for its active site (residues 10-37). This lid contains a cleavage site between S23 and N24 that is an important locus of EcA2 cleavage by lysosomal proteases, responsible for the *in vivo* inactivation of the enzyme [38]. N24 is located at the N-terminal flexible loop, which in Aginasa has reduced flexibility in the presence of L-Asp bound to the active site, causing it to remain in its closed conformation. The increased protease accessibility of N24 and, consequently, the higher propensity for proteolysis demonstrated by Leuginase is most likely related to the open-loop conformation observed for the EcA2 present in this product. With the addition of L-Asp, Leuginase exhibits a degradation profile similar to Aginasa, indicating that L-Asp binding exerts a protective effect against proteolysis. We did not detect any additional protection against proteolysis after incubation with L-Glu. These observations confirm the results obtained by NMR, demonstrating the inefficiency of L-Glu in stabilizing the EcA2 closed-lid conformation.

The measurement of the distances between L-Asp and the residues that constitute the catalytic site of EcA2 (Table S5), along with the STD-NMR data, revealed that L-Asp H^α^ is in close contact with H^γ21^ of V27, located at the N-terminal loop of EcA2 (Fig. 11 and 12). L-Glu causes no more than 50% of the conformational effect in the loop, likely reflecting the much lower affinity of this amino acid for EcA2 (1.9 x 10^−5^ M for L-Asn vs. 5.9 x 10^−3^ M for L-Gln) [36]. In addition to its direct role in ligand binding stabilization, Val27 is also in close contact with Gly11, part of the N-terminal hinge of the loop that allows this element to transit between conformations upon ligand binding [12], limiting hinge mobility. Contacts of Val27 with the loop residue Tyr25 were observed and possibly contributes with the restriction of Tyr25 sidechain flexibility, as observed by NMR. Moreover, as identified in the crystal structure, the residue Val27 interacts with Gly57, which is part of the active site (Table S6).

**Figure 12.**
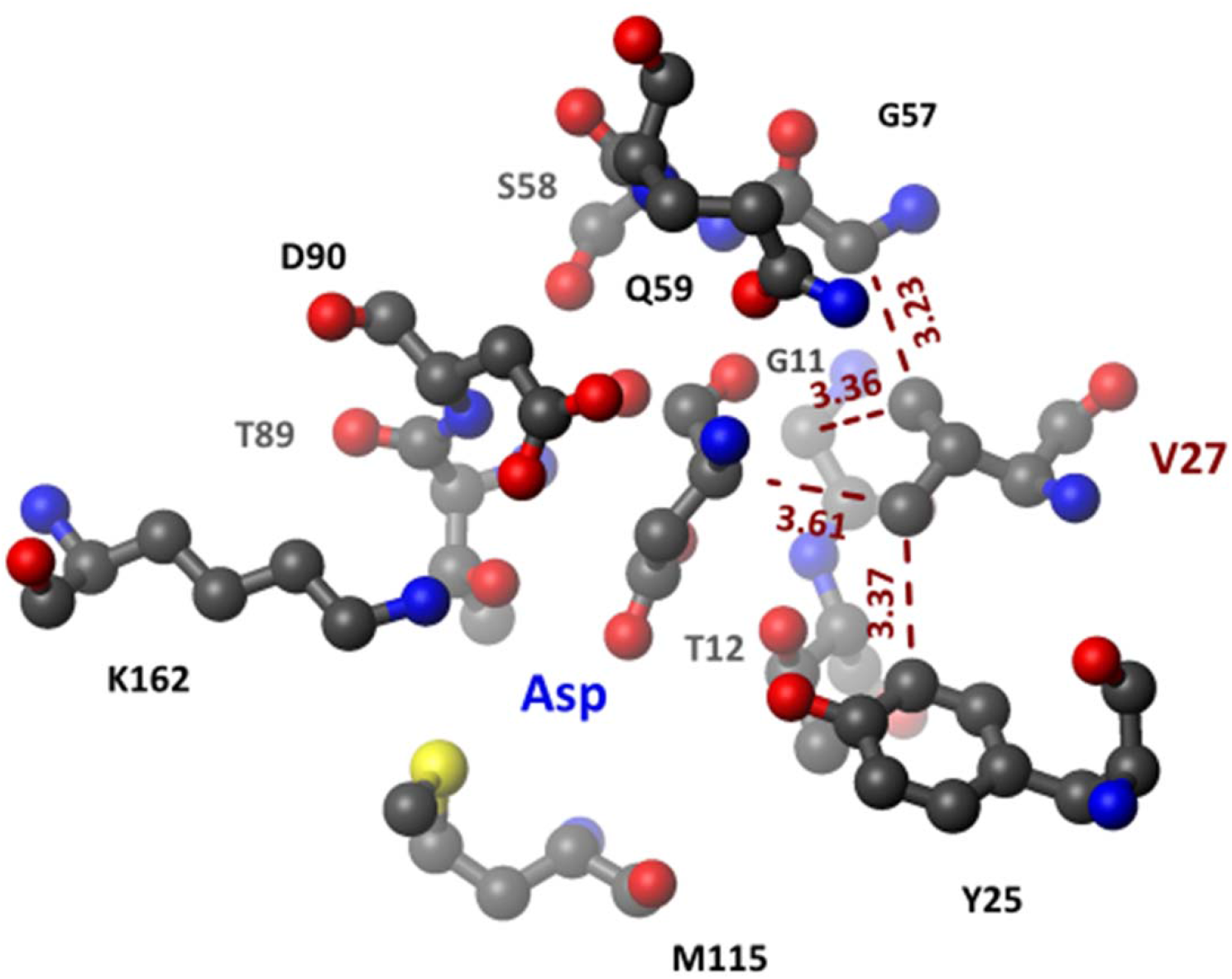
Stereochemical analysis of EcA2 catalytic site upon L-Asp binding. The figure illustrates the ball and stick representation of the EcA2 catalytic site (Leuginase: Asp – PDB Id.: 6UOG) pointing out Val27 interactions (represented by red dashed lines) together with the distances in Angstroms estimated between carbon atoms. All the estimated distances between Val27 and the catalytic site residues and the ligand L-Asp can be found in Table S5 and S6. Despite ligand binding being maintained mostly by hydrogen bonds (LigPlot shown in Fig. S11), Val27 appears to play a role linking the ligand binding with conformational change toward the active closed lid form. As demonstrated in the figure, Val27 presents a network of hydrophobic interactions, with the ligand L-Asp (H^α^), active site residues G57 and G11, and the sidechain of the loop residue Y25.

Sequence alignment of different species of L-asparaginase using BLASTp (31,32) was highly conserved for the residues involved in binding and catalysis (Fig. 13). Interestingly, Val27 is present only in EcA2 and L-asparaginase from *Saccharomyces cerevisiae*, while other L-asparaginases have alanine or serine at this position. A previous study using EcA2 variants with mutations on V27 replaced with the apolar amino acids Met and Leu showed a decrease in catalytic activity for L-Asn and L-Gln to about 20% of the wild type activity due to a reduced K_cat_ value.[41]. Another study carried out with engineered ErA2 variants with mutations in A31, the corresponding residue to EcA2 V27 in this isoform, demonstrated substantial variations in substrate affinity and L-asparaginase/L-glutaminase activity [5]. The crystallographic structure of ErA2 A31I, a variant with higher affinity for both L-Asn and L-Gln, revealed that this mutation forced a different conformation in the N-terminal flexible loop in comparison to the wild-type enzyme. However, the authors could not explain the decreased *K*_M_ value observed for this mutant since no obvious conclusion could be made from the structure and they speculated that this mutation influenced the dynamics of the loop clorsure and ultimately the substrate affinity.

**Figure 13.**
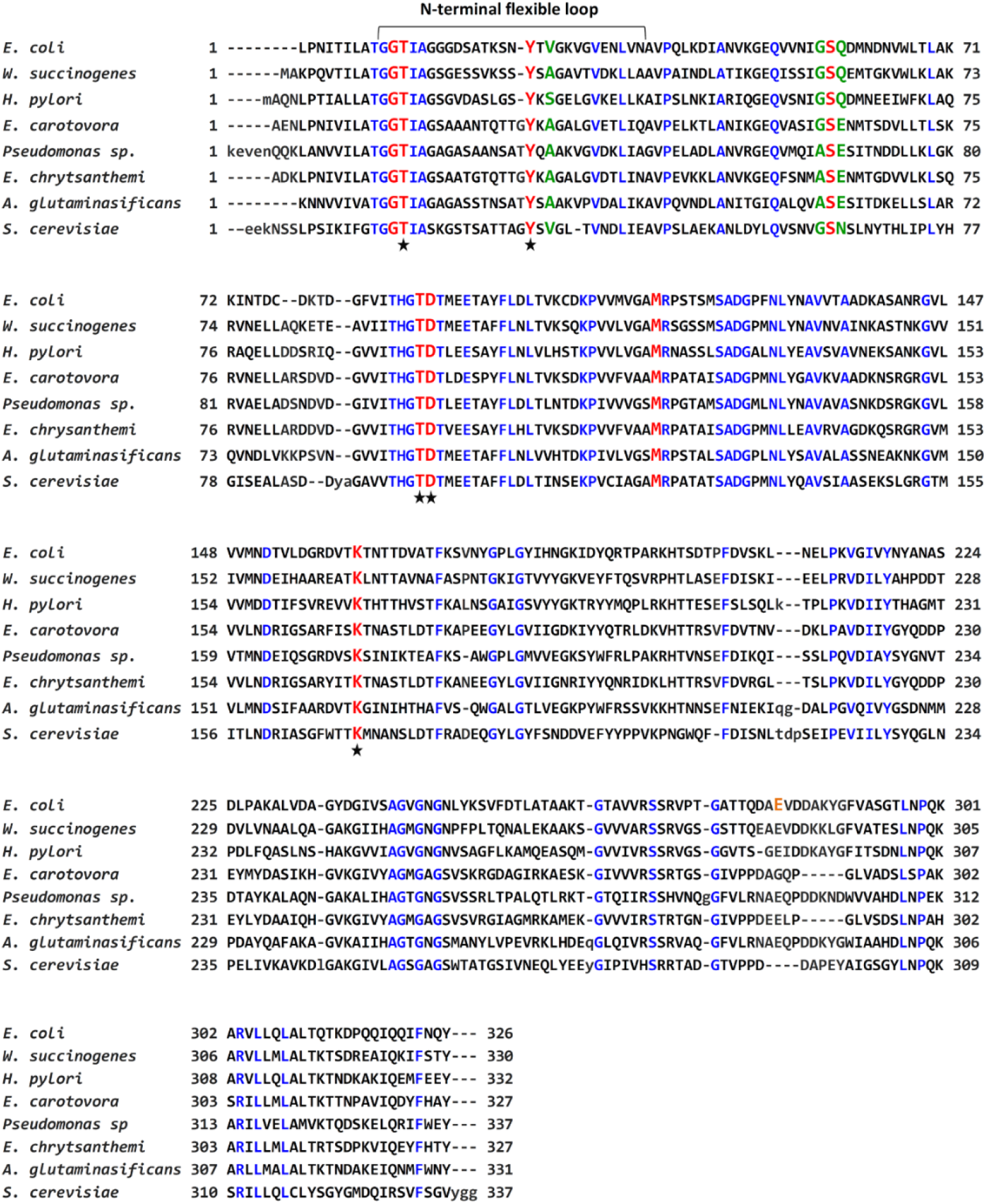
Sequence alignment of L-asparaginases. Conserved residues are colored in blue. Conserved active site residues are highlighted in red, with those described as being directly involved in catalysis indicated by black stars. Residues within the catalytic site that are not well-conserved are shown in green. Residue E283, which shows high Cα RMS in Leuginase Apo structure (Figure 8B), is highlighted in orange. This difference is due to its role in ligand binding when the residue is in the adjacent chain that forms the intimate dimer (Fig. S11). The flexible N-terminal loop boundaries of EcA2, identified through the Cα RMS displacement analysis, are indicated in the figure. The UniProt entries of the sequences used in this alignment, with identities (I) and homology (H) compared to EcA2 indicated in parenthesis, were: *Escherichia coli* P00805, *Wolinella succinogenes* (I – 55%; H – 72%) P50286, *Helicobacter pylori* (I – 50%; H – 69%) Q9ZLB9, *Erwinia carotovora* (I – 49%; H – 65%) Q7WWK9, *Pseudomonas sp*. (I – 47%; H – 67%) P10182, *Erwinia chrysanthemi* P06608 (I – 48%; H – 63%), *Acinetobacter glutaminasificans* (I – 45%; H – 64%) P10172, and *Saccharomyces cerevisiae* (I – 41%; H – 56%) P0CX77. The sequence alignment was performed using BLASTp.

The superposition of the L-asparaginase catalytic site structure shows that stereochemical features are also highly conserved, and the main difference lies in the orientation of the loop forming the lid over the catalytic site (Fig. 14). This is an intrinsic characteristic that affects the catalytic activity and substrate specificity of these enzymes, with EcA2 having one with the lowest *K*_M_ values of the enzymes studied (34,35,36), despite the similarities in sequence and overall folding.

**Figure 14.**
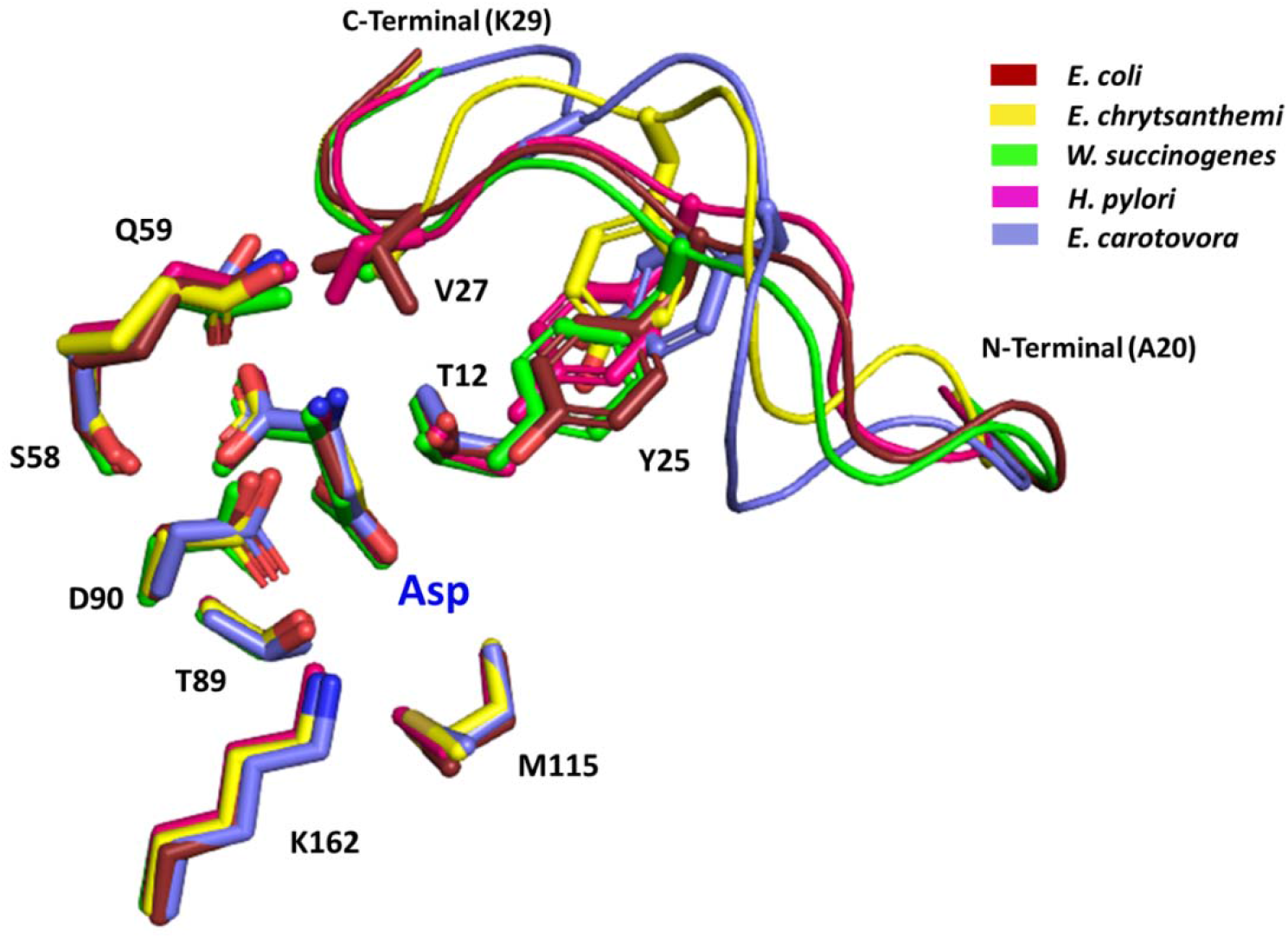
Superposition of the L-asparaginase catalytic site. The structural alignment demonstrates that L-asparaginases from different species exhibit high conservation of catalytic site structural features with a major difference located at the loop forming the lid coordinates, specially Val27 and Tyr25, which are crucial for loop stabilization. Only L-asparaginases with structures solved with L-Asp were superposed. Residues are labeled according to the EcA2 sequence. The PDB entries of corresponding structures used for this alignment were *Wolinella succinogenes* PDB ID 5K3O, *Escherichia coli* (Leuginase-Asp) PDB ID 6UOG, *Erwinia chrysanthemi* PDB ID 5F52, *Erwinia carotovora* PDB ID 2GVN, *Helicobacter pylori* PDB ID 2WLT.

These observations indicate that Val27 plays a crucial role in the conformational changes in the lid following substrate binding and stabilization of the catalytic site conformation. This could explain the differences in catalytic activity, substrate affinity, and loop conformation exhibited for different L-asparaginases. This residue is then considered a good target for enzyme engineering.

### 4.2. Formulation aspects

We found that both Aginasa and Leuginase had molecular weights that are similar and compatible with the mature form of EcA2 (Uniprot P00805; amino acids 23-348), with an average mass of 34.593,94 Da and monoisotopic mass of 34.572,56 Da. We performed MS analysis of both products under two entirely different setups and spectrometers and elucidated their crystal structures from each dataset. All data showed redundant evidence for mature EcA2 as the major component of the formulation. Previous work reported molecular weight of 36.8 kDa for Leuginase, which would correspond to the full amino acid sequence of EcA2, including the 22-amino acid long-signal peptide [18]. However, data from tandem-mass spectrometry sequencing from this same study did not find the complete signal peptide. We have no reasonable explanation for these conflicting results except for the fact that we used different batches. However, it seems implausible that the production method and the resulting biopharmaceutical product would have changed within such a short interval (the difference between batches was approximately one year). Unfortunately, the manufacturing processes of synthetic and biosynthetic pharmaceutical products are proprietary information, which is only made available to regulatory agencies in documents such as the Drug Master Files (DMFs) or New Drug Applications (NDAs) of the U.S. Food and Drug Administration (FDA).

Interestingly, we found that the EcA2 in Leuginase has an average mass of 15 Da less than expected. This difference between Leuginase and Aginasa suggests the presence of different amino acid sequences that were not detected within the coverage limits of the mass spectrometry proteolytic fragmentation analysis. Although some set of amino acid substitutions could result in a 15 Da difference, it would be counterproductive to engage in conjecture. Given that both products show similar activity *in vitro* (10,11,25) and similar physicochemical and ultra-structural properties as described in this study, we assume that this difference may represent a molecular signature to identify the compound, although we cannot confirm whether this is intentional or unintentional.

The product leaflets do not typically disclose the components of the formulation; non-guided analysis of the product often results in such components being found. For example, L-Asp has been found in the crystallographic structures of the native EcA2 formulation Elspar [7] and the Y25F variant of EcA2 [46]. Since neither compound was incorporated as soaking/co-crystallization, the L-Asp was part of the product formulation. We found both mannitol and PEG in Leuginase, which could be present due to purification by crystallization [47] or to increase the stability of the formulation. The product leaflet, however, does not disclose any adjuvants in the formulation. Furthermore, we found aspartate in Aginasa, which is also not disclosed in its leaflet. All these compounds are justifiable from a downstream perspective, including during final product stabilization. However, the lack of full disclosure of their composition and the implications for their *in vitro* activity/physicochemical behavior hamper the development of a pharmacopeic monograph, which is only currently available for EcA2 in the Chinese Pharmacopoeia (Office TS 2017,), with no mention of monitoring for related substances including PEG, mannitol, aspartate, sulfate or any other stabilizer or residual component of the downstream process. Moreover, both products – Aginasa and Leuginase – entered the Brazilian health care system through an international purchase conducted by the Ministry of Health. In addition, no bidding process was required for the use of this drug in Brazil. Based on our findings, we suggest that a drug master file must be made available to regulatory agencies, along with clinical data and full disclosure of the components in the pharmaceutical product leaflet. Doing so would provide greater awareness of the products being used and would ensure fuller transparency to the clinicians and health care system.

### 4.3. Further therapeutic aspects

EcA2 has long been used as a therapeutic agent with impressive results in the success rate of ALL treatment (37,38). However, some patients are not responsive to this therapy, perhaps owing to the inactivation of the enzyme by immune response or the proteolytic degradation of EcA2 [51]. It is worth noting that this erratic response to therapy is also observed with the use of PEGylated EcA2 [38]. In this context, the development of other L-asparaginases with higher *in vivo* stability against inactivation and higher catalytic efficiency may benefit from the conformational and dynamic features unveiled in this work. However, this is not a complete mapping of the conformational and stability landscape of EcA2. Further studies are advised with an extended mapping of the interaction with other amino acids, as well as long-term stability studies in the presence of the amino acids selected as the best candidates for stabilization (most likely L-Asp).

Despite the similar conformational and stability profile of Aginasa and Leuginase in the presence of L-Asp as shown by IMS, crystal structure, NMR measurements, and partial proteolysis, we are still intrigued by the 15 Da difference between them. This shall merit further detailed investigation since, as far as we are aware, no drug master file has been disclosed. It remains to be determined whether this difference is related to the subtle, but significant, differences in their thermostability and hydrodynamic radii.

Our results demonstrate that the selectivity of L-asparaginase is correlated with the stabilization of its active site lid conformation. In light of our results and the accumulated evidence from the literature, the selectivity of L-Asp in binding and its effect on the dynamics of the active site lid support its use as a kinetic and thermodynamic stabilizing agent of this enzyme.

In addition, the pharmacopeial monograph for EcA2 in the Chinese Pharmacopoeia (Office TS 2017), determine specifications assays such as identification by high performance liquid chromatography, pH, purity assessed by size exclusion chromatography, and enzymatic activity to determine potency in vitro. However, these techniques provide information that corresponds to a global average of the protein structure and chemical identity and subsequently are usually uncapable to detect changes in amino acid sequence and specific conformational changes. Therefore, a more detailed physical-chemical characterization and structural investigation, using analytical techniques with higher accuracy and resolution such as NMR and MS of the final product is imperative to avoid harsh clinical disparities.

## Supporting information

Supportng Material

## Data availability

The crystallographic structure factors and pdb models presented in this paper have been deposited in the Protein Data Bank (PDB) with the following codes: 6UOD, 6UOG, 6UOH. All other datasets generated during and/or analyzed during the current study are available from the corresponding author on reasonable request.

## Acknowledgments

We acknowledge Dr. Rafael Andrade and Jessica Moreira for their technical support on protein biochemistry at Plataforma Avançada de Biomoléculas/UFRJ. We would like to thank LNLS-CNPEM (MX2 and SAS1 beamlines), UEMP-UFRJ, CENABIO-UFRJ, CEMBIO-UFRJ, LDRX-IF-UFF and LAMAC-DIMAV-INMETRO and their staff for the access to their analytical facilities. This research used resources of the Brazilian Synchrotron Light Laboratory (LNLS), an open national facility operated by the Brazilian Centre for Research in Energy and Materials (CNPEM) for the Brazilian Ministry for Science, Technology, Innovations and Communications (MCTIC).

## Funding and additional information

This study was supported by Fundação Carlos Chagas Filho de Amparo à Pesquisa do Estado do Rio de Janeiro (grants E-26/202.998/2017-BOLSA, to LMTRL and E-26/290.076/2017 to MSA), by the Conselho Nacional de Desenvolvimento Científico (grant PQ2/311582/2017-6, to LMTRL), by the Coordenação de Aperfeiçoamento de Pessoal de Nível Superior (CAPES, Finance Code #001) and the Programa Nacional de Apoio ao Desenvolvimento da Metrologia, Qualidade e Tecnologia (PRONAMETRO, to LMTRL) from the Instituto Nacional de Metrologia, Qualidade e Tecnologia (INMETRO). The funding agencies had no role in the study design, data collection and analysis, or decision to publish or prepare of the manuscript.

## References

[1] S. Patil, J. Coutsouvelis, A. Spencer, Asparaginase in the management of adult acute lymphoblastic leukaemia: Is it used appropriately?, Cancer Treat. Rev. 37 (2011) 202–207. https://doi.org/10.1016/j.ctrv.2010.08.002.

[2] Y. Tabe, p.L. Lorenzi, M. Konopleva, Amino acid metabolism in hematologic malignancies and the era of targeted therapy, Blood. 134 (2019) 1014–1023. https://doi.org/10.1182/blood.2019001034.

[3] Glutaminase activity determines cytotoxicity of L-asparaginases on most leukemia cell lines, Leuk. Res. 39 (2015) 757–762. https://doi.org/10.1016/j.leukres.2015.04.008.

[4] T.N. Seyfried, G. Yu, J.C. Maroon, D.P.D. Agostino, Press-pulseL: a novel therapeutic strategy for the metabolic management of cancer, Nutr. Metab. (Lond). 14:19 (2017) 1–17. https://doi.org/10.1186/s12986-017-0178-2.

[5] H.A. Nguyen, Y. Su, A. Lavie, Design and characterization of erwinia chrysanthemi L-asparaginase variants with diminished L-glutaminase activity, J. Biol. Chem. 291 (2016) 17664–17676. https://doi.org/10.1074/jbc.M116.728485.

[6] R. Gribskov, A left-handed crossover involved in amidohydrolase structure catalysis of Erwinia chrysanthemi L-asparaginase with bound L-aspartate, 328 (1993) 275–279.

[7] Crystal structure of Escherichia coli L-asparaginase, an enzyme used in cancer therapy, Proc. Natl. Acad. Sci. U. S. A. 90 (1993) 1474–1478.

[8] N.E. Labrou, A.C. Papageorgiou, V.I. Avramis, Structure-Function Relationships Asparaginases and Clinical Applications of L-Asparaginases, Curr. Med. Chem. 17 (2010) 2183–2195.

[9] M. Maggi, S.D. Mittelman, J.H. Parmentier, G. Colombo, M. Meli, J.M. Whitmire, D.S. Merrell, J. Whitelegge, C. Scotti, A protease-resistant Escherichia coli asparaginase with outstanding stability and enhanced antileukaemic activity in vitro, Sci. Rep. 7 (2017) 1–16. https://doi.org/10.1038/s41598-017-15075-4.

[10] J. Aghaiypour, K; Wlodawer,A; Lubkowski, Structural Basis for the Activity and Substrate Specificity of, Biochemistry. 40 (2001) 5655–5664. https://doi.org/10.1074/jbc.M110.107177.

[11] Structural Insight into Substrate Selectivity of Erwinia chrysanthemi L-asparaginase, Biochemistry. 55 (2016) 1246–1253. https://doi.org/10.1021/acs.biochem.5b01351.

[12] H.A. Nguyen, D.L. Durden, A. Lavie, The differential ability of asparagine and glutamine in promoting the closed/active enzyme conformation rationalizes the Wolinella succinogenes L-asparaginase substrate specificity, Sci. Rep. 7 (2017) 1–14. https://doi.org/10.1038/srep41643.

[13] Development of L-Asparaginase Biobetters: Current Research Status and Review of the Desirable Quality Profiles, Front. Bioeng. Biotechnol. 6 (2019). https://doi.org/10.3389/fbioe.2018.00212.

[14] B. Asselin, C. Rizzari, Asparaginase pharmacokinetics and implications of therapeutic drug monitoring, Leuk. Lymphoma. 56 (2015) 2273–2280. https://doi.org/10.3109/10428194.2014.1003056.

[15] M.K. Schwartz, E.D. Lash, H.F. Oettgen, F.A. Tomao, L-asparaginase activity in plasma and other biological fluids, Cancer. 25 (1970) 244–252. https://doi.org/10.1002/1097-0142(197002)25:2<244::AID-CNCR2820250203>3.0.CO;2-V.

[16] J. Boos, G. Werber, E. Ahlke, p. Schulze-Westhoff, U. Nowak-Göttl, G. Würthwein, E.J. Verspohl, J. Ritter, H. Jürgens, Monitoring of asparaginase activity and asparagine levels in children on different asparaginase preparations, Eur. J. Cancer Part A. 32 (1996) 1544–1550. https://doi.org/10.1016/0959-8049(96)00131-1.

[17] Monitoring asparaginase activity in middle-income countries, Lancet. Oncol. 19 (2018) 1149–1150. https://doi.org/10.1016/S1470-2045(18)30584-9.

[18] Low Bioavailability and High Immunogenicity of a New Brand of E. coli l-Asparaginase with Active Host Contaminating Proteins, EBioMedicine. 30 (2018) 158–166. https://doi.org/10.1016/j.ebiom.2018.03.005.

[19] M.K. Parr, O. Montacir, H. Montacir, Physicochemical characterization of biopharmaceuticals, J. Pharm. Biomed. Anal. 130 (2016) 366–389. https://doi.org/10.1016/j.jpba.2016.05.028.

[20] mMass data miner: an open source alternative for mass spectrometric data analysis, Rapid Commun. Mass Spectrom. RCM. 22 (2008) 905–908. https://doi.org/10.1002/rcm.3444.

[21] L.C. Palmieri, B. Melo-ferreira, C.A. Braga, G.N. Fontes, L.J. Mattos, L.M.T.R. Lima, Stepwise oligomerization ofmurine amylin and assembly ofamyloid fibrils., Biophys. Chem. 180–181 (2013) 135–144. https://doi.org/10.1016/j.bpc.2013.07.013.

[22] Use of ion mobility mass spectrometry and a collision cross-section algorithm to study an organometallic ruthenium anticancer complex and its adducts with a DNA oligonucleotide, Rapid Commun. Mass Spectrom. RCM. 23 (2009) 3563–3569. https://doi.org/10.1002/rcm.4285.

[23] W. Kabsch, research papers XDS research papers, Acta Crystallogr. Sect. D Biol. Crystallogr. D66 (2010) 125–132. https://doi.org/10.1107/S0907444909047337.

[24] Coot: model-building tools for molecular graphics, Acta Crystallogr. Sect. D Biol. Crystallogr. 60 (2004) 2126–2132. https://doi.org/10.1107/S0907444904019158.

[25] REFMAC5 for the refinement of macromolecular crystal structures, Acta Crystallogr. Sect. D Biol. Crystallogr. 67 (2011) 355–367. https://doi.org/10.1107/S0907444911001314.

[26] The Phenix Software for Automated Determination of Macromolecular Structures, Methods. 55 (2011) 94. https://doi.org/10.1016/j.ymeth.2011.07.005.

[27] A. Viegas, J. Manso, F.L. Nobrega, E.J. Cabrita, Saturation-transfer difference (STD) NMR: A simple and fast method for ligand screening and characterization of protein binding, J. Chem. Educ. 88 (2011) 990–994. https://doi.org/10.1021/ed101169t.

[28] S.M. Kelly, T.J. Jess, N.C. Price, How to study proteins by circular dichroism, 1751 (2005) 119–139. https://doi.org/10.1016/j.bbapap.2005.06.005.

[29] The Small-Angle X-ray Scattering Beamline of the Brazilian Synchrotron Light Laboratory, J. Appl. Crystallogr. 30 (1997) 880–883. https://doi.org/10.1107/S0021889897001829.

[30] FIT2D: a multi-purpose data reduction, analysis and visualization program, J. Appl. Crystallogr. 49 (2016) 646–652. https://doi.org/10.1107/S1600576716000455.

[31] ATSAS 2.8: a comprehensive data analysis suite for small-angle scattering from macromolecular solutions, J. Appl. Crystallogr. 50 (2017) 1212–1225. https://doi.org/10.1107/S1600576717007786.

[32] P. V Konarev, V. V Volkov, A. V Sokolova, H.J. Koch, D.I. Svergun, PRIMUSL: a Windows PC-based system for small-angle scattering data analysis, J. Appl. Cryst. 36 (2003) 1277–1282.

[33] B.T. Ruotolo, J.L.P. Benesch, A.M. Sandercock, S. Hyung, C. V Robinson, Ion mobility – mass spectrometry analysis of large protein complexes, 3 (2008) 1139–1152. https://doi.org/10.1038/nprot.2008.78.

[34] R. Salbo, M.F. Bush, H. Naver, I. Campuzano, C. V Robinson, I. Pettersson, T.J.D. Jørgensen, K.F. Haselmann, Traveling-wave ion mobility mass spectrometry of protein complexes□: accurate calibrated collision cross-sections of human insulin oligomers, Rapid Commun. Mass Spectrom. 26 (2012) 1181–1193. https://doi.org/10.1002/rcm.6211.

[35] C. Derst, A. Wehner, V. Specht, K. -H Röhm, States and Functions of Tyrosine Residues in Escherichia coli Asparaginase II, Eur. J. Biochem. 224 (1994) 533–540. https://doi.org/10.1111/j.1432-1033.1994.00533.x.

[36] N. Nakamura, Y. Morikawa, M. Tanaka, L-asparaginase from escherichia coli: Part II. Substrate specificity studies, Agric. Biol. Chem. 35 (1971) 743– 751. https://doi.org/10.1080/00021369.1971.10859979.

[37] M. Sanches, S. Krauchenco, I. Polikarpov, Structure, Substrate Complexation and Reaction Mechanism of Bacterial Asparaginases, Curr. Chem. Biol. 1 (2012) 75–86. https://doi.org/10.2174/2212796810701010075.

[38] A dyad of lymphoblastic lysosomal cysteine proteases degrades the antileukemic drug L-asparaginase, J. Clin. Invest. 119 (2009) 1964–1973. https://doi.org/10.1172/JCI37977.

[39] S.F. Altschul, T.L. Madden, A.A. Schäffer, J. Zhang, Z. Zhang, W. Miller, D.J. Lipman, Gapped BLAST and PSI-BLAST□: a new generation of protein database search programs, 25 (1997) 3389–3402.

[40] S.F. Altschul, J.C. Wootton, E.M. Gertz, R. Agarwala, A. Morgulis, A.A. Scha, Protein database searches using compositionally adjusted substitution matrices, 272 (2005) 5101–5109. https://doi.org/10.1111/j.1742-4658.2005.04945.x.

[41] Engineering the substrate specificity of Escherichia coli asparaginase. II. Selective reduction of glutaminase activity by amino acid replacements at position 248, Protein Sci. A Publ. Protein Soc. 9 (2000) 2009–2017. https://doi.org/10.1110/ps.9.10.2009.

[42] A.P. Sudhir, B.R. Dave, A.S. Prajapati, K. Panchal, D. Patel, R.B. Subramanian, Characterization of a Recombinant Glutaminase-Free l-Asparaginase (ansA3) Enzyme with High Catalytic Activity from Bacillus licheniformis, Appl. Biochem. Biotechnol. 174 (2014) 2504–2515. https://doi.org/10.1007/s12010-014-1200-z.

[43] A. Beckett, D. Gervais, What makes a good new therapeutic l l1 asparaginase□?, World J. Microbiol. Biotechnol. 9 (2019) 1–13. https://doi.org/10.1007/s11274-019-2731-9.

[44] J.J.M. Cachumba, F.A.F. Antunes, G.F.D. Peres, L.P. Brumano, J.C. Dos Santos, S.S. Da Silva, Current applications and different approaches for microbial L-asparaginase production, Brazilian J. Microbiol. 47 (2016) 77–85. https://doi.org/10.1016/j.bjm.2016.10.004.

[45] L.M.T.R. Lima, T.S. Araujo, F.C.L. Almeida, M.S. Almeida, Monitoring, Lancet Oncol. 19 (2018) e574. https://doi.org/10.1016/S1470-2045(18)30656-9.

[46] Structures of two highly homologous bacterial l--asparaginases: a case of enantiomorphic space groups, Acta Crystallogr. Sect. D Biol. Crystallogr. 57 (2001) 369–377. https://doi.org/10.1107/S0907444900020175.

[47] X. Yu, Y. Wu, F. Huang, J. Ulrich, J. Wang, Purification of Recombinant L-asparaginase II Using Solvent Freeze Out (SFO) Technology, Chem. Eng. Technol. 41 (2018) 1080–1085. https://doi.org/10.1002/ceat.201700569.

[48] Office TS 2017, Pharmacopoeia of the People’s Republic of China 2015., in: B, Beijing: The Stationery Office., n.d.

[49] R.A. Egler, S.P. Ahuja, Y. Matloub, L-asparaginase in the treatment of patients with acute lymphoblastic leukemia, J. Pharmacol. Pharmacother. 7 (2016) 62–71. https://doi.org/10.4103/0976-500X.184769.

[50] M. Mohseni, H. Uludag, J.M. Brandwein, Advances in biology of acute lymphoblastic leukemia (ALL) and therapeutic implications., Am. J. Blood Res. 8 (2018) 29–56.

[51] C. Lanvers-Kaminsky, Asparaginase pharmacology: Challenges still to be faced, Cancer Chemother. Pharmacol. 79 (2017) 439–450. https://doi.org/10.1007/s00280-016-3236-y

