## Supplementary material for "Biophysical characterization of two commercially available preparations of the drug containing *Escherichia coli* L-Asparaginase 2": Supportng Material

**Dynamic modulation of the *Escherichia coli* L-Asparaginase 2 conformational landscape by ligand binding: formulation and therapeutic implications**

Talita Stelling de Araújo^1,2^, Sandra M. N. Scapin^4^, William de Andrade^3^, Maira Fasciotti^4^, Mariana T. Q. de Magalhães^5^, Marcius S. Almeida*^1,2^, Luís Maurício T. R. Lima*^3,4^

**List of contents:**

**Table S1.** Melting transition temperatures (Tm) and slope of Aginasa and Leuginase calculated from CD thermal denaturation curves at 220 nm.

**Table S2.** Aginasa, Leuginase, and Leuginase: Asp retention time and respective hydrodynamic radius estimated by interpolation from curves.

**Table S3.** Data collection and refinement statistics.

**Table S4.** Secondary-structure alignment of monomers from the Aginasa, apo Leuginase, and Leuginase: L-Asp.

**Table S5.** Secondary-structure alignment of monomers from the Aginasa, apo Leuginase, and Leuginase: L-Asp.

**Table S6.** Secondary-structure alignment of monomers from the Aginasa, apo Leuginase, and Leuginase: L-Asp.

**Fig. S1.** Enzymatic digestion of Aginasa and Leuginase with trypsin and peptide identification by MALDI-TOF/TOF mass spectrometry.

**Fig. S2.** Co-polymer comprising asparaginase formulation as revealed by LC-ESI-MS.

**Fig. S3.** Quantitative analysis of Aginasa and Leuginase formulation constituents by NMR.

**Fig. S4.** Asparaginase structural models used in the analysis of oligomeric distribution in solution from SAXS data

**Fig. S5.** Oligomer analysis of EcA2 structural models with Aginasa scattering curves.

**Fig. S6.** TSKgel G3000SW_XL_ column standard curve.

**Fig. S7.** Size-exclusion chromatography analysis of Aginasa, Leuginase, and Leuginase supplemented with 1 mM L-Asp.

**Fig. S8.** Mapping interaction of amino acid with EcA2 by MS.

**Fig. S9.** Multiple displays of 1D ^1^H NMR spectra of Aginasa and Leuginase whole formulation and Leuginase added with 1 mM of L-Asp.

**Fig. S10.** Characterization of limited proteolysis EcA2 products by LC-MS.

**Fig. S11.** LigPlot diagram of EcA2 catalytic site upon L-Asp binding.

**Table S1.** Melting transition temperatures (Tm) and slope of Aginasa and Leuginase calculated from CD thermal denaturation curves at 220 nm.

|  | **Melting Temperature (°C)** | **Slope** |
| --- | --- | --- |
| **Aginasa** | 67.52 ± 0.10 | 0.21 ± 0.01 |
| **Aginasa + L-Asp** | 66.78 ± 0.13 | 0.22 ± 0.01 |
| **LeugiNase** | 67.87 ± 0.08 | 0.39 ± 0.02 |
| **LeugiNase + L-Asp** | 67.91 ± 0.06 | 0.44 ± 0.02 |

**Table S2.** Aginasa, Leuginase, and Leuginase: Asp retention time and respective hydrodynamic radius estimated by interpolation from curves.

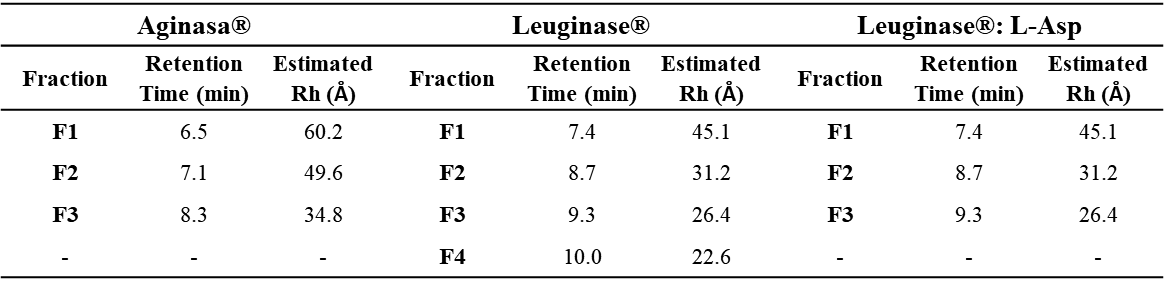

**Table S3.** Data collection and refinement statistics.

|  | **holo (L-ASP bound)** | | | **Apo** |
| --- | --- | --- | --- | --- |
| **Protein preparation** | | **Aginasa** | **LeugiNase: L-ASP** | **LeugiNase** |
| **PDB Code** | | **6UOH** | **6UOG** | **6UOD** |
| **Data collection** | | | | |
| Space group | | C121 | P 1 21 1 | C121 |
| **Unit cell** | |  |  |  |
| a; b; c (Å) | | 75.38; 133.55; 63.91 | 139.99; 60.16; 151.11 | 152.22; 60.08; 142.79 |
| α; β; γ (°) | | 90.00; 110.344; 90.00 | 90.00; 117.31; 90.00 | 90.00; 117.99; 90.00 |
| Wavelength (Å) | | 1.46 | 1.46 | 1.46 |
| Resolution (Å) | | 44.60 – 1.89 (1.89 – 2.00) | 47.07 – 2.1 (2.1 – 2.26) | 47.11 – 2.31 (2.31 – 2.45) |
| CC_1/2_ | | 0.999 (0.85) | 0.993 (0.762) | 0.998 (0.967) |
| R_merge_ | | 0.04 (0.45) | 0.08 (0.358) | 0.03 (0.14) |
| Average I/σ (I) | | 18.09 (1.88) | 7.660 (2.09) | 13.85 (3.53) |
| Completeness (%) | | 77.2 (62.7) | 87.4 (65.0) | 73.0 (76.5) |
| Wilson B-factors (Å^2^) | | 38.136 | 39.599 | 41.218 |
| **Data Processing and Refinement** | | |  |  |
| Refinement program | | REFIMAC5/PHENIX | REFIMAC5 | REFIMAC5/PHENIX |
| Resolution (Å) | | 32.02 – 2.10 (2.1 – 2.13) | 47.07 – 2.29 (2.29 – 2.35) | 38.58 - 2.4 (2.40 – 2.44) |
| Completeness (%) | | 96.3 (91.0) | 95.9 (71.4) | 92.7 (89.0) |
| Number of Reflections | | 65715 (2857) | 92730 (5037) | 80962 (3502) |
| R_work_ | | 0.222 (0.296) | 0.200 (0.216) | 0.198 (0.254) |
| R_free_ | | 0.277 (0.332) | 0.234 (0.271) | 0.252 (0.326) |
| Average B-factors (Å^2^) | | 39.0 | 36.253 | 36.0 |
| Protein molecules per asymmetric unit | | 2 | 8 | 4 |
| Asp molecules | | 2 | 8 | 0 |
| Water molecules | | 362 | 615 | 479 |
| **R.m.s. deviation from ideal:** | | |  |  |
| Bond length (Å) | | 0.007 | 0.002 | 0.008 |
| Bond angles (°) | | 0.865 | 1.250 | 0.919 |
| **Ramachandran plot statistics (%)** | | |  |  |
| Most favored | | 97.07 | 97.8 | 96.08 |
| Additionally allowed | | 2.47 | 2.2 | 3.92 |
| Outlier | | 0.46 | 0 | 0 |

High resolution shell in parenthesis; r.m.s., root-mean square.

**Table S4.** Secondary-structure alignment of monomers from the Aginasa, apo Leuginase, and Leuginase: L-Asp. Root-mean square values were obtained with ProSMART (CCP4 suite).

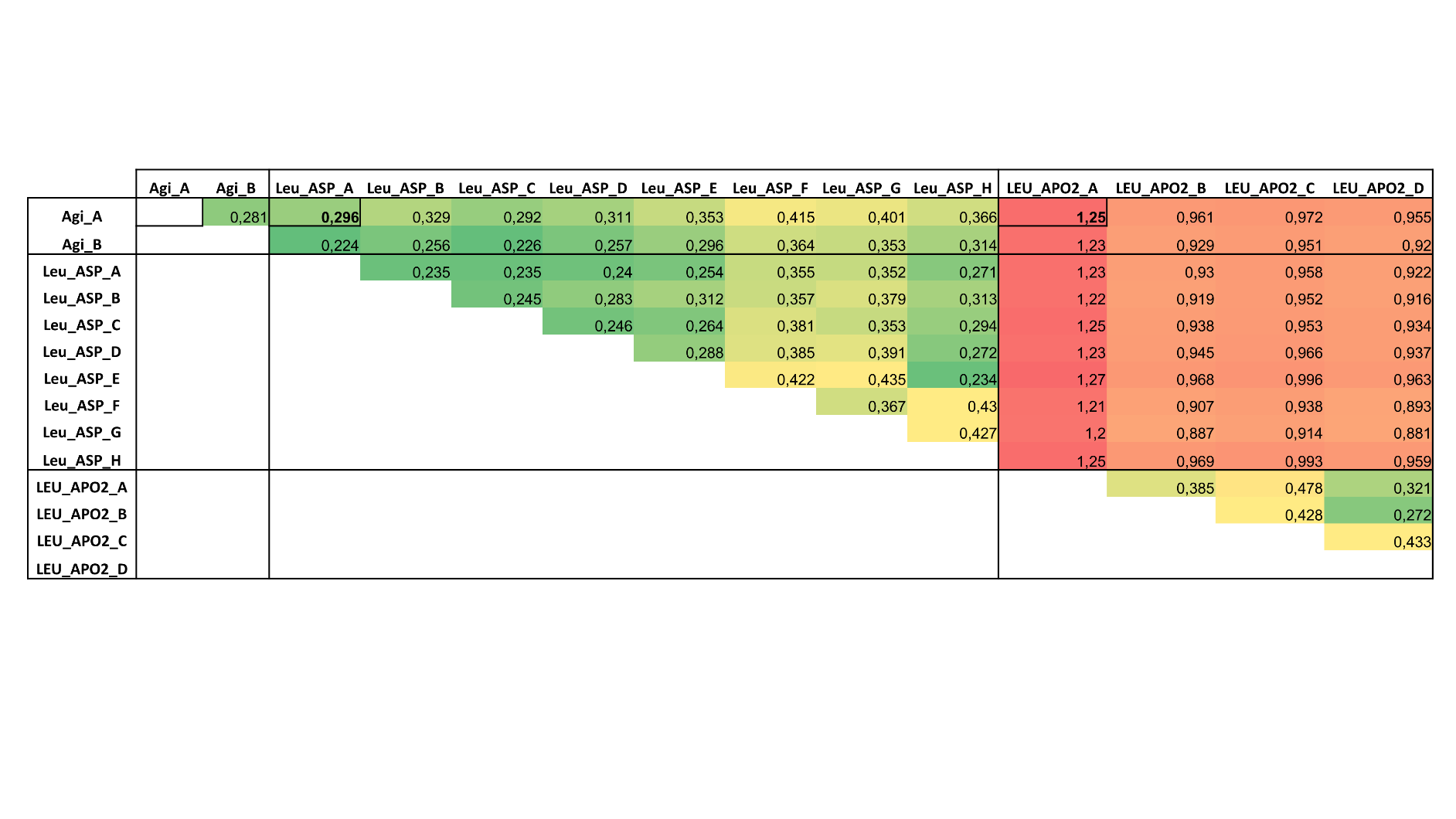

**Table S5.** Measurement of the distances between Val27 and the residues that constitute the catalytic site of EcA2. The distances between carbons was estimated with MolMol with cut-off of 5 Å.

| **Val27**  **Atom** | **Catalytic Site Residue/Atom** | | | **Distance (Å)** |
| --- | --- | --- | --- | --- |
| CG1 | 57 | GLY | CA | 3.23 |
| CG1 | 11 | GLY | CA | 3.36 |
| CG2 | 25 | TYR | CE2 | 3.37 |
| CG2 | 12 | THR | CA | 4.02 |
| CG2 | 25 | TYR | CD2 | 4.05 |
| CB | 57 | GLY | CA | 4.21 |
| CG2 | 12 | THR | CB | 4.29 |
| CG2 | 25 | TYR | CZ | 4.34 |
| CB | 11 | GLY | CA | 4.60 |
| CG1 | 12 | THR | CA | 4.65 |
| CB | 25 | TYR | CE2 | 4.65 |
| CA | 25 | TYR | CD2 | 4.71 |
| CB | 12 | THR | CA | 4.72 |
| CA | 25 | TYR | CE2 | 4.75 |
| CG2 | 11 | GLY | CA | 4.81 |
| CA | 12 | THR | CA | 4.87 |

**Table S6.** Measurement of the distances between L-Asp and the residues that constitute the catalytic site of EcA2. The distances between heavy atoms were obtained with MolMol with cut-off of 5 Å.

| **L-Asp Atom** | **Catalytic Site Residue/Atom** | | | **Distance (Å)** |  | **L-Asp Atom** | **Catalytic Site Residue/Atom** | | | **Distance (Å)** |
| --- | --- | --- | --- | --- | --- | --- | --- | --- | --- | --- |
| OXT | 58 | SER | OG | 2.34 |  | OD2 | 115 | MET | CA | 4.28 |
| CG | 12 | THR | OG1 | 2.62 |  | OD2 | 115 | MET | S | 4.28 |
| OD2 | 89 | THR | OG1 | 2.65 |  | CG | 90 | ASP | OD1 | 4.29 |
| N | 90 | ASP | OD2 | 2.75 |  | OXT | 89 | THR | OG1 | 4.29 |
| N | 59 | GLN | OE1 | 2.77 |  | C | 57 | GLY | C | 4.31 |
| O | 58 | SER | N | 2.80 |  | CB | 90 | ASP | N | 4.32 |
| OD1 | 12 | THR | OG1 | 2.81 |  | CA | **27** | **VAL** | **CG1** | 4.33 |
| OD2 | 12 | THR | OG1 | 2.86 |  | CB | 12 | THR | CB | 4.34 |
| CG | 89 | THR | OG1 | 2.93 |  | O | 27 | VAL | CB | 4.35 |
| OD1 | 12 | THR | N | 2.94 |  | CG | 25 | TYR | OH | 4.35 |
| OXT | 90 | ASP | N | 2.94 |  | OXT | 58 | SER | C | 4.36 |
| OD1 | 89 | THR | N | 3.05 |  | N | 90 | ASP | CB | 4.36 |
| OXT | 89 | THR | N | 3.13 |  | O | 89 | THR | N | 4.36 |
| CB | 12 | THR | OG1 | 3.17 |  | CA | 12 | THR | N | 4.41 |
| OXT | 90 | ASP | CB | 3.18 |  | C | **27** | **VAL** | **CG2** | 4.42 |
| OXT | 90 | ASP | CG | 3.25 |  | OD2 | 162 | LYS+ | NZ | 4.42 |
| O | 57 | GLY | CA | 3.28 |  | OD2 | 12 | THR | N | 4.43 |
| CB | 90 | ASP | OD1 | 3.28 |  | C | 12 | THR | OG1 | 4.43 |
| C | 58 | SER | OG | 3.29 |  | OD2 | 89 | THR | CA | 4.46 |
| OD1 | 12 | THR | CB | 3.30 |  | C | 12 | THR | N | 4.48 |
| OD2 | 12 | THR | CB | 3.34 |  | O | 59 | GLN | N | 4.48 |
| O | 11 | GLY | CA | 3.37 |  | OD2 | 12 | THR | CA | 4.50 |
| O | **27** | **VAL** | **CG1** | 3.38 |  | C | 59 | GLN | N | 4.50 |
| CG | 12 | THR | CB | 3.39 |  | CA | **27** | **VAL** | **CB** | 4.51 |
| N | 90 | ASP | CG | 3.40 |  | CG | 90 | ASP | N | 4.52 |
| OXT | 58 | SER | CB | 3.42 |  | CB | **27** | **VAL** | **CG2** | 4.55 |
| C | 59 | GLN | OE1 | 3.44 |  | CG | 12 | THR | CG2 | 4.55 |
| CA | 12 | THR | OG1 | 3.47 |  | O | 57 | GLY | N | 4.56 |
| O | 57 | GLY | C | 3.48 |  | N | 12 | THR | OG1 | 4.57 |
| O | 59 | GLN | OE1 | 3.48 |  | CB | 90 | ASP | CB | 4.58 |
| O | 58 | SER | OG | 3.48 |  | CA | 90 | ASP | CB | 4.60 |
| N | 59 | GLN | CD | 3.49 |  | CA | 25 | TYR | OH | 4.60 |
| C | 58 | SER | N | 3.51 |  | C | 59 | GLN | CD | 4.62 |
| CB | 89 | THR | OG1 | 3.51 |  | CB | 162 | LYS+ | NZ | 4.62 |
| OD1 | 89 | THR | OG1 | 3.51 |  | CA | 59 | GLN | CD | 4.64 |
| CG | 89 | THR | N | 3.53 |  | OXT | 57 | GLY | C | 4.65 |
| OD2 | 89 | THR | CG2 | 3.56 |  | CA | 89 | THR | N | 4.65 |
| OXT | 90 | ASP | OD2 | 3.60 |  | OD2 | 115 | MET | O | 4.65 |
| CA | **27** | **VAL** | **CG2** | 3.61 |  | O | 59 | GLN | CD | 4.66 |
| OXT | 58 | SER | N | 3.61 |  | OD1 | 90 | ASP | N | 4.66 |
| OXT | 90 | ASP | CA | 3.63 |  | CA | 58 | SER | OG | 4.66 |
| CB | 90 | ASP | CG | 3.63 |  | O | 58 | SER | C | 4.67 |
| CA | 90 | ASP | OD2 | 3.64 |  | CB | 89 | THR | CB | 4.67 |
| OD2 | 89 | THR | CB | 3.65 |  | OD2 | 90 | ASP | OD1 | 4.67 |
| OXT | 90 | ASP | OD1 | 3.66 |  | CB | 12 | THR | N | 4.68 |
| CA | 59 | GLN | OE1 | 3.67 |  | OXT | 90 | ASP | C | 4.70 |
| OD1 | 12 | THR | CA | 3.71 |  | OD1 | 12 | THR | CG2 | 4.70 |
| N | 59 | GLN | NE2 | 3.75 |  | O | 57 | GLY | O | 4.71 |
| CB | 90 | ASP | OD2 | 3.80 |  | CA | 12 | THR | CB | 4.72 |
| CA | 90 | ASP | CG | 3.82 |  | OD2 | 115 | MET | N | 4.72 |
| CG | 12 | THR | N | 3.82 |  | N | 59 | GLN | CG | 4.73 |
| N | 90 | ASP | OD1 | 3.82 |  | OD2 | 115 | MET | C | 4.75 |
| O | 58 | SER | CB | 3.83 |  | O | 12 | THR | OG1 | 4.76 |
| O | 58 | SER | CA | 3.84 |  | OD1 | 12 | THR | C | 4.77 |
| C | 89 | THR | N | 3.84 |  | C | 90 | ASP | CA | 4.77 |
| OD1 | 11 | GLY | C | 3.87 |  | N | **27** | **VAL** | **CB** | 4.77 |
| OD1 | 11 | GLY | CA | 3.87 |  | CA | 90 | ASP | N | 4.77 |
| CG | 89 | THR | CB | 3.88 |  | CA | 89 | THR | OG1 | 4.77 |
| OXT | 59 | GLN | OE1 | 3.88 |  | CA | 58 | SER | N | 4.79 |
| C | 90 | ASP | CG | 3.89 |  | CG | 90 | ASP | CG | 4.80 |
| C | 90 | ASP | OD2 | 3.91 |  | OD1 | **27** | **VAL** | **CG2** | 4.80 |
| OXT | 58 | SER | CA | 3.94 |  | C | 11 | GLY | C | 4.81 |
| OXT | 89 | THR | C | 3.95 |  | CG | 162 | LYS+ | NZ | 4.81 |
| N | **27** | **VAL** | **CG2** | 3.95 |  | C | 58 | SER | C | 4.82 |
| OD1 | 89 | THR | CG2 | 3.95 |  | N | **27** | **VAL** | **CG1** | 4.82 |
| CA | 90 | ASP | OD1 | 3.96 |  | N | 59 | GLN | CB | 4.84 |
| CG | 89 | THR | CG2 | 3.98 |  | C | 89 | THR | OG1 | 4.84 |
| OD1 | 89 | THR | CB | 4.06 |  | OXT | 89 | THR | CB | 4.85 |
| C | 58 | SER | CB | 4.07 |  | CB | 89 | THR | CA | 4.87 |
| OD2 | 89 | THR | N | 4.07 |  | CG | 11 | GLY | C | 4.89 |
| CB | 25 | TYR | OH | 4.07 |  | N | 59 | GLN | N | 4.89 |
| C | 90 | ASP | N | 4.10 |  | CG | 89 | THR | C | 4.90 |
| OXT | 89 | THR | CA | 4.11 |  | N | 25 | TYR | OH | 4.90 |
| O | 11 | GLY | N | 4.12 |  | CA | 11 | GLY | CA | 4.91 |
| OD1 | 89 | THR | CA | 4.13 |  | CG | 27 | VAL | CG2 | 4.91 |
| C | 90 | ASP | CB | 4.14 |  | O | 90 | ASP | OD2 | 4.92 |
| O | 11 | GLY | C | 4.15 |  | C | 27 | VAL | CB | 4.93 |
| O | 12 | THR | N | 4.16 |  | N | 58 | SER | N | 4.93 |
| OD2 | 25 | TYR | OH | 4.18 |  | OXT | 11 | GLY | CA | 4.95 |
| O | **27** | **VAL** | **CG2** | **4.20** |  | CA | 59 | GLN | NE2 | 4.95 |
| CB | 89 | THR | N | 4.20 |  | OD1 | 89 | THR | C | 4.96 |
| OD2 | 12 | THR | CG2 | 4.21 |  | N | 57 | GLY | CA | 4.96 |
| OXT | 59 | GLN | N | 4.21 |  | OXT | 57 | GLY | CA | 4.96 |
| C | 11 | GLY | CA | 4.23 |  | C | 89 | THR | CA | 4.98 |
| CG | 12 | THR | CA | 4.23 |  | CB | 90 | ASP | CA | 4.98 |
| C | 57 | GLY | CA | 4.25 |  | CA | 57 | GLY | CA | 4.99 |
| CG | 89 | THR | CA | 4.26 |  | O | 90 | ASP | CG | 4.99 |
| C | **27** | **VAL** | **CG1** | 4.26 |  |  |  |  |  |  |
| C | 90 | ASP | OD1 | 4.27 |  |  |  |  |  |  |
| C | 58 | SER | CA | 4.28 |  |  |  |  |  |  |

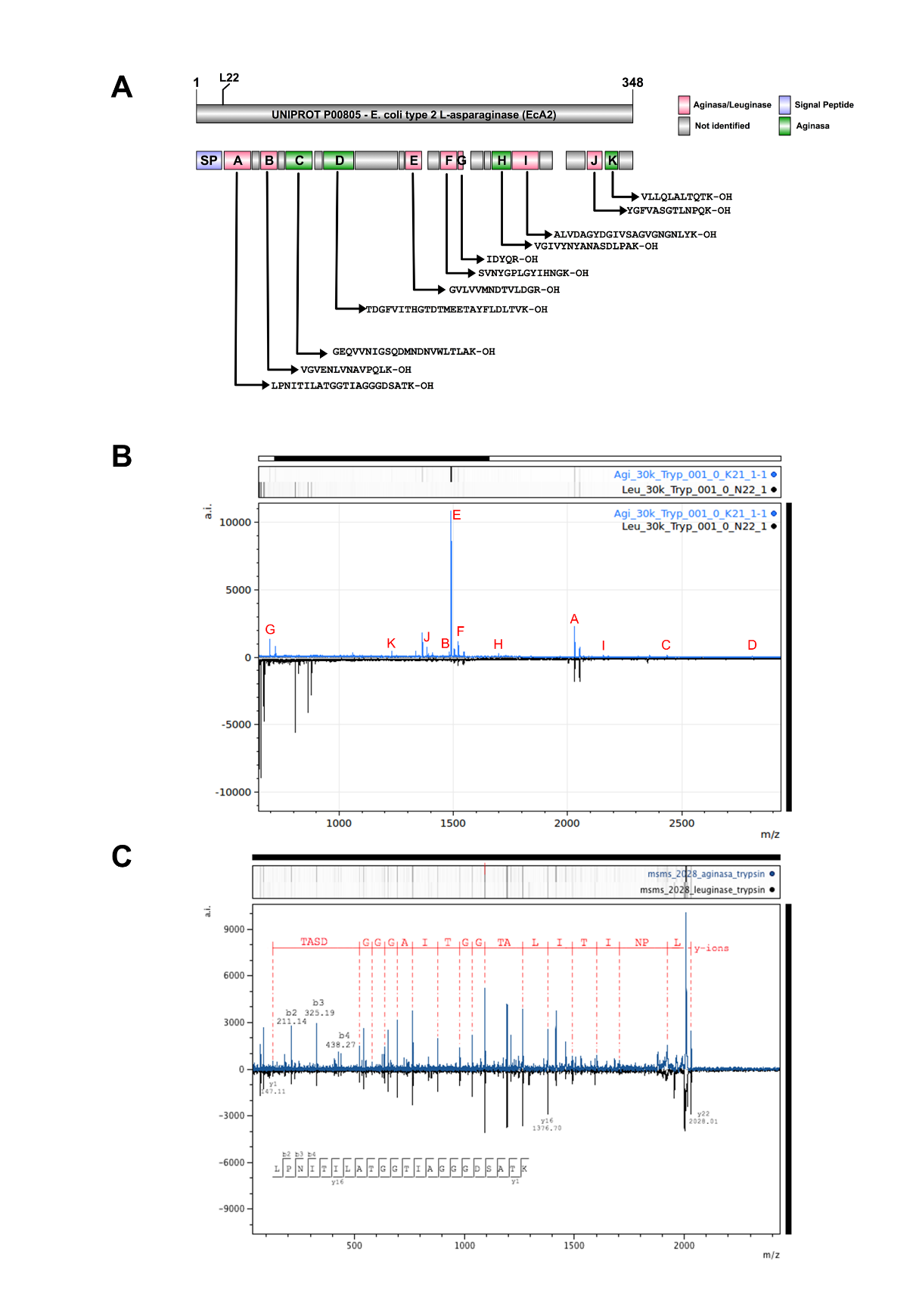

**Fig. S1.** Enzymatic digestion of Aginasa and Leuginase with trypsin and peptide identification by MALDI-TOF/TOF mass spectrometry. **A.** Schematic representation EcA2 amino acid sequence tryptic digestion coverage for both Aginasa and Leuginase proteins **B.** The molecular masses comparison of all tryptic fragments for Aginasa and Leuginase tryptic evaluated by MALDI-TOF/MS. **C.** MS/MS spectra assignment for Aginasa and Leuginase [M+H]^+^ 2028.09 m/z . The peptides were fragmented by MALDI TOF MS/MS experiments showing the same profile. The resulting data were interpreted manually using mMass software reveling a peptide with primary sequence LPNITILATGGTIAGGGDSATK-OH corresponding to the N-terminal 22 residues fragment.

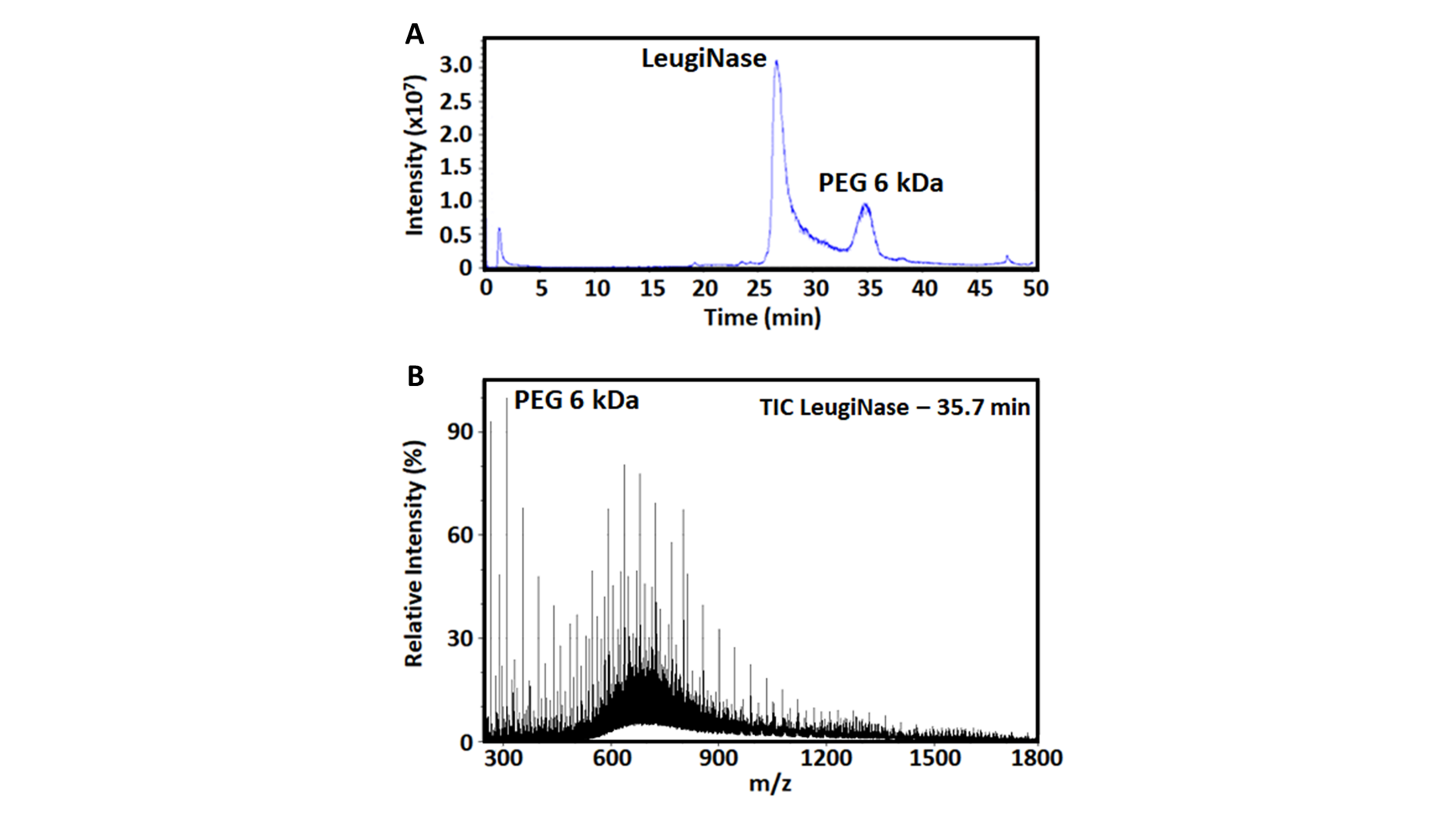

**Fig. S2.** Co-polymer comprising asparaginase formulation as revealed by LC-ESI-MS. **A.** The total ion current (TIC) chromatogram of Leuginase showed a major peak corresponding to the intact protein and a peak corresponding to PEG. **B.** The deconvoluted spectra of the 34.7 min peak revealed the presence of PEG 6000 in Leuginase formulation.

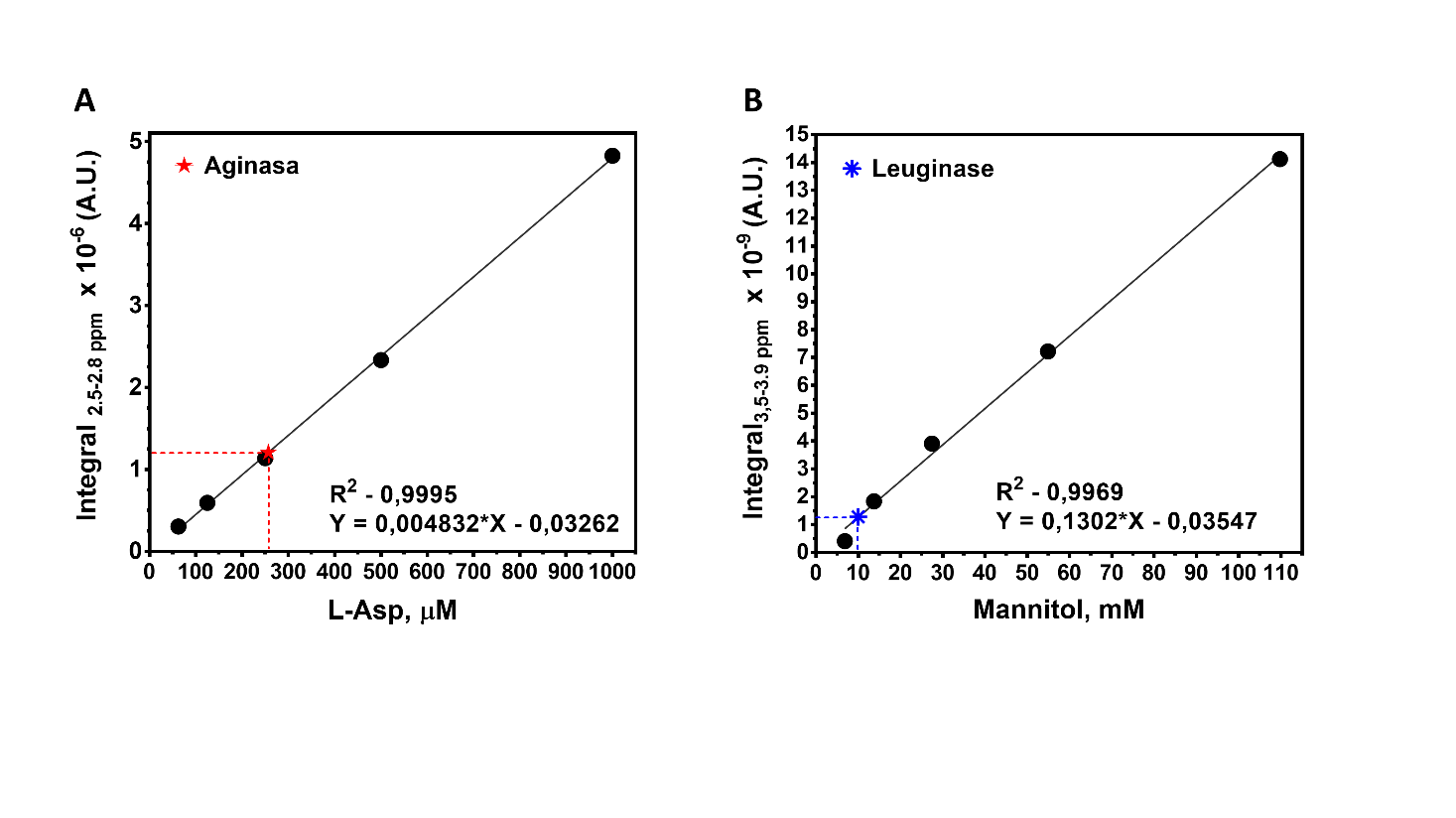

**Fig. S3.** Quantitative analysis of Aginasa and Leuginase formulation constituents by NMR. Quantitative analysis shows the presence of approximately **A.** 260 µM L-ASP in Aginasa and **B.** 10 mM of Mannitol in Leuginase formulation. Integrals from 1D ^1^H spectra acquired for standard solutions of the formulation constituents identified were used to fit a standard linear curve. The concentration of the constituents was obtained by interpolation of the curve.

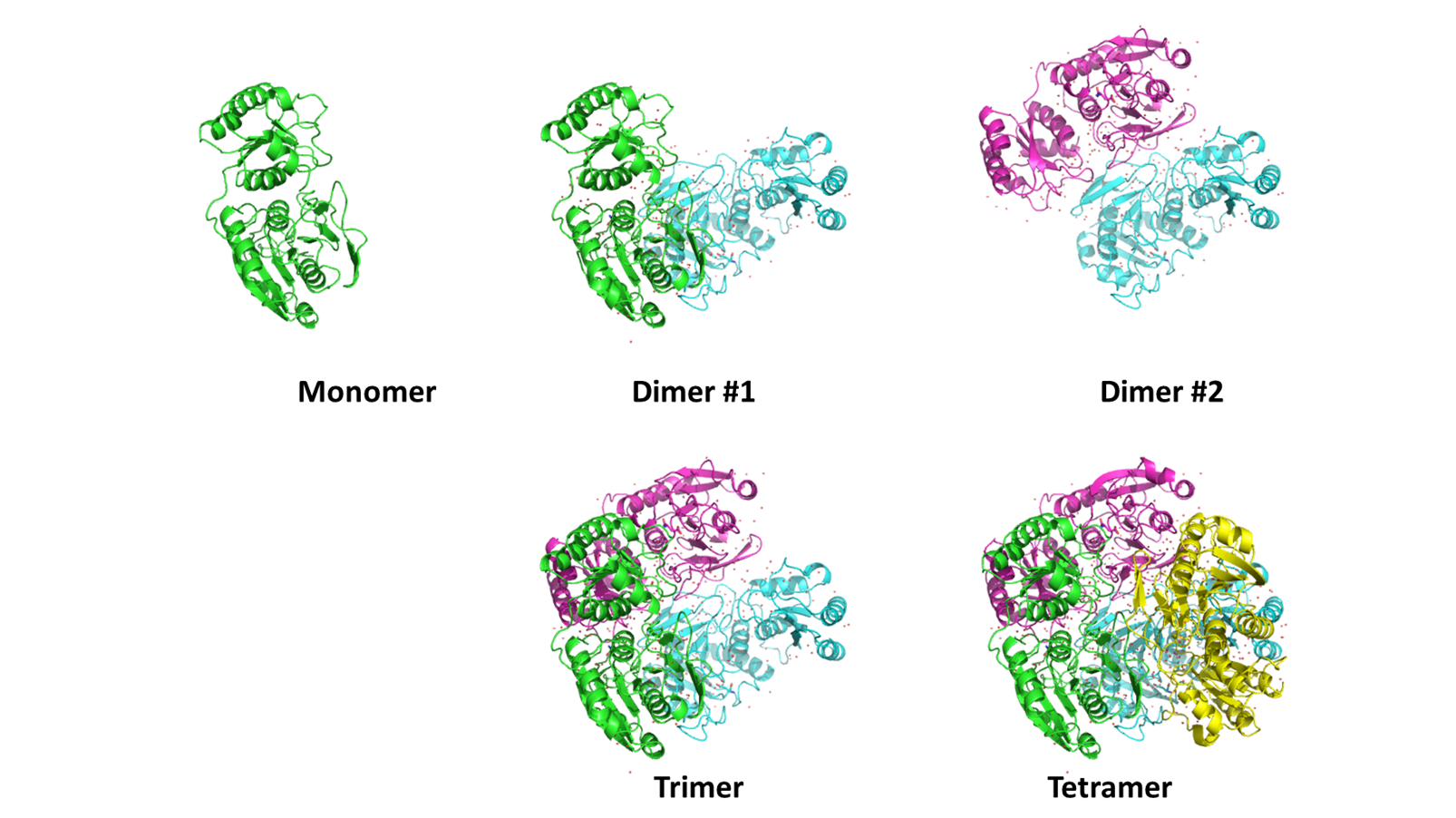

**Fig. S4.** Asparaginase structural models used in the analysis of oligomeric distribution in solution from SAXS data. Models were generated with PyMOL using crystal structure model 3ECA.pdb.

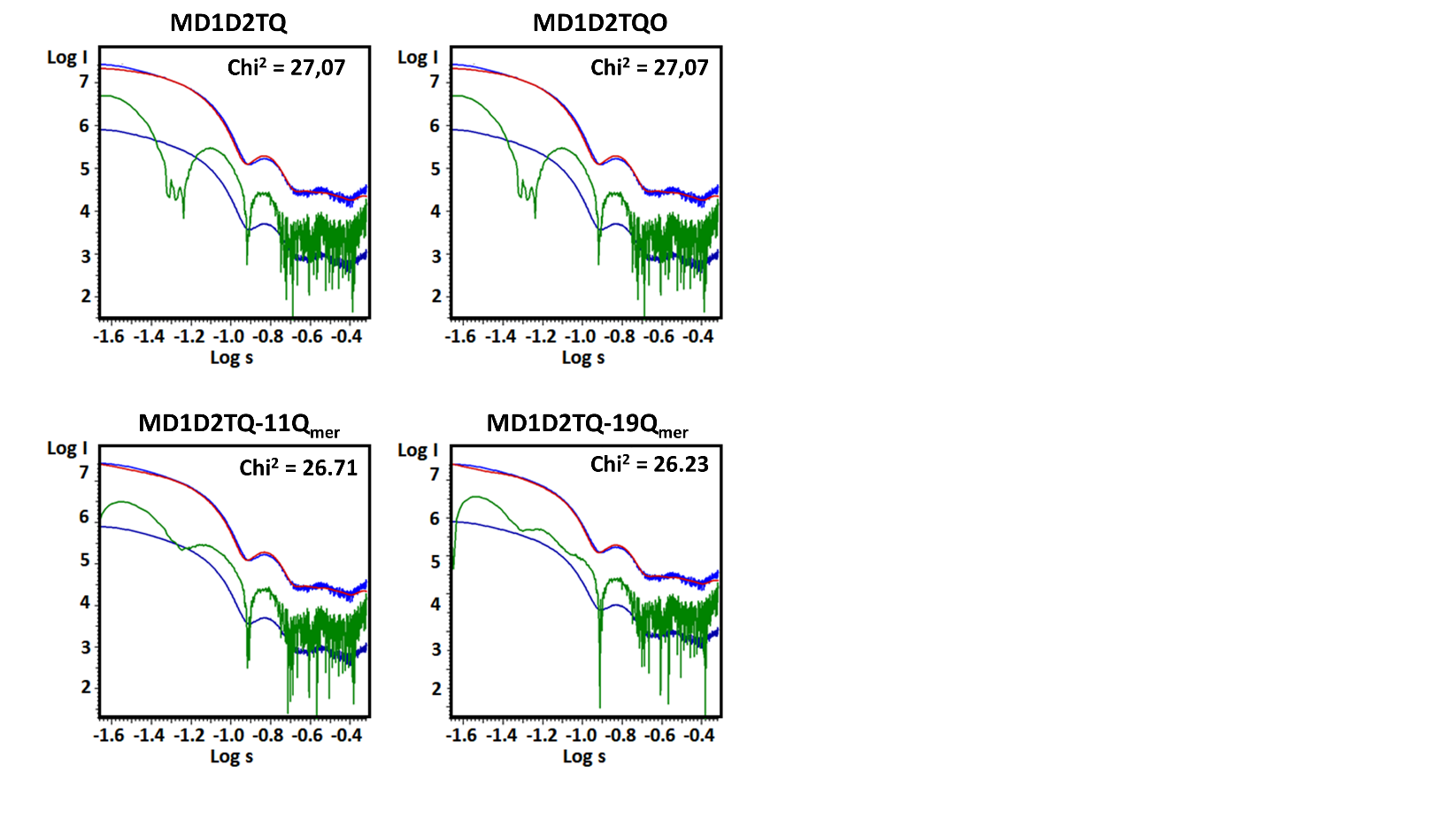

**Fig. S5.** Oligomer analysis of EcA2 structural models with Aginasa scattering curves. Curves were adjusted with models comprising monomer (M), dimers (D1, D2), Trimer (T), tetramer (Q), octamer (O), and higher-order oligomer (11mer or 19-mer, as generated from 3ECA with PyMOL). Curves are raw data (light blue), best adjustment with model (red), residuals (green), and adjustment error (dark blue).

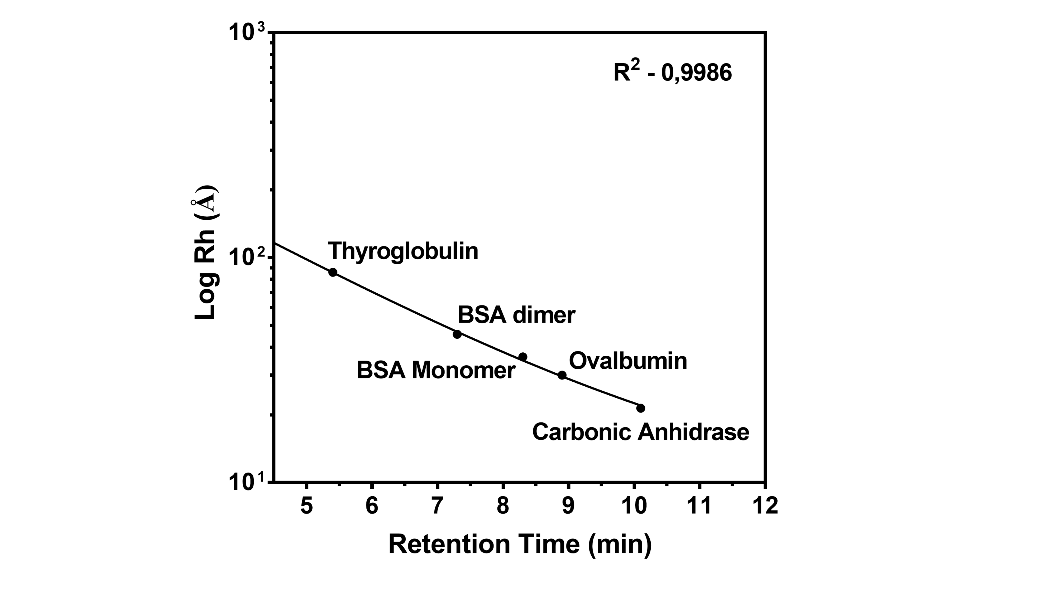

**Fig. S6.** TSKgel G3000SW_XL_ column standard curve. The elution times were plotted against the hydrodynamic radius of each standard protein and fitted to an exponential one-phase decay function. The standard globular proteins used and their hydrodynamic radius were thyroglobulin – 86 Å, bovine serum albumin – dimer 45.6 Å and monomer 36.2 Å, Ovalbumin– 30.5 Å, and carbonic anhydrase – 21.4 Å.

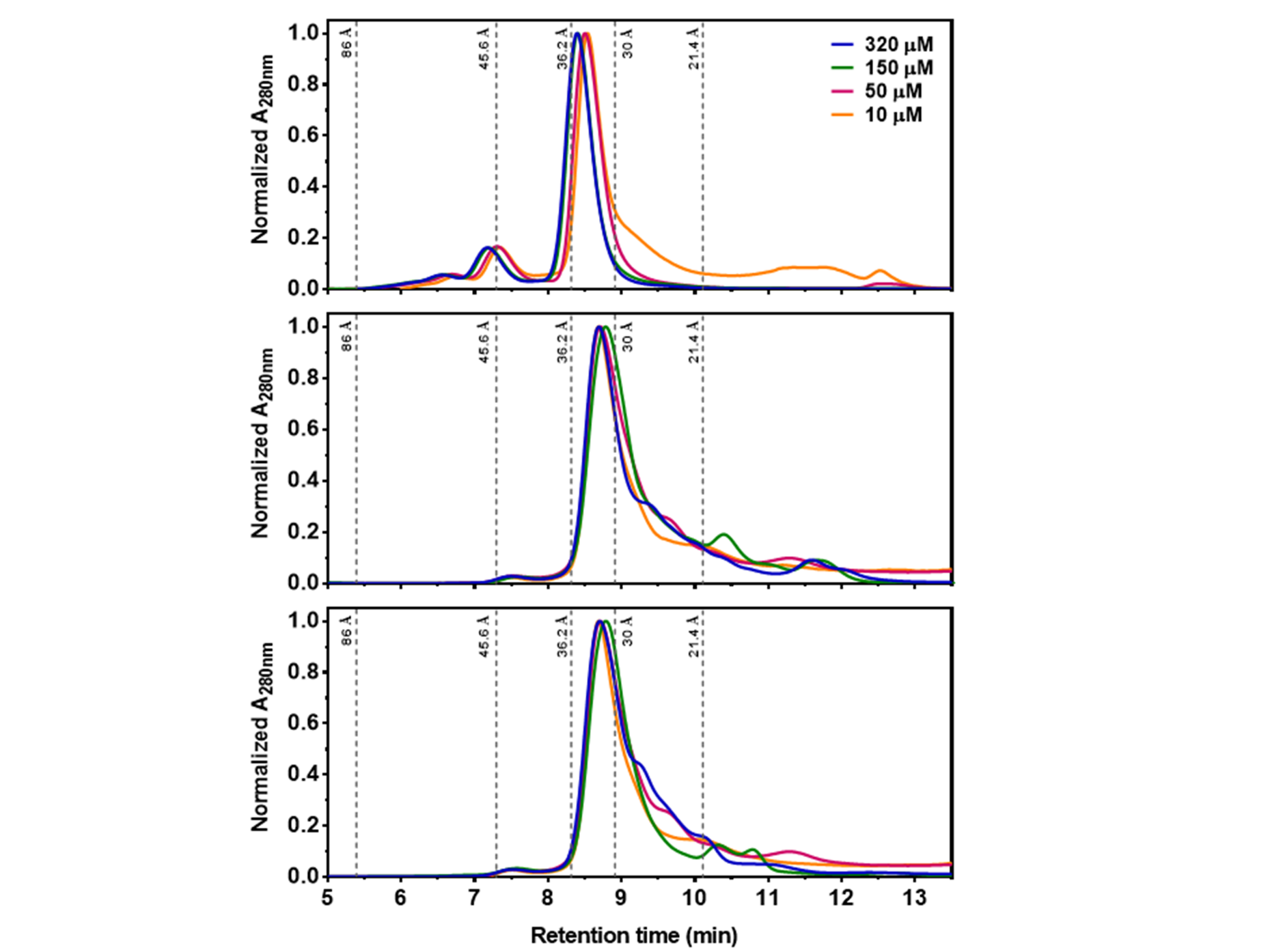

**A**

**C**

**B**

**Fig. S7.** Size-exclusion chromatography analysis of Aginasa (A), Leuginase (B), and Leuginase supplemented with 1 mM L-Asp (C). The hydrodynamic radius of standard globular proteins is indicated at their retention volume. Chromatograms in blue represent the sample at 320 µM, in green 320 µM, pink 50 µM and in yellow 10 µM. The absorbance at 280 nm was normalized to facilitate comparison of chromatographic runs.

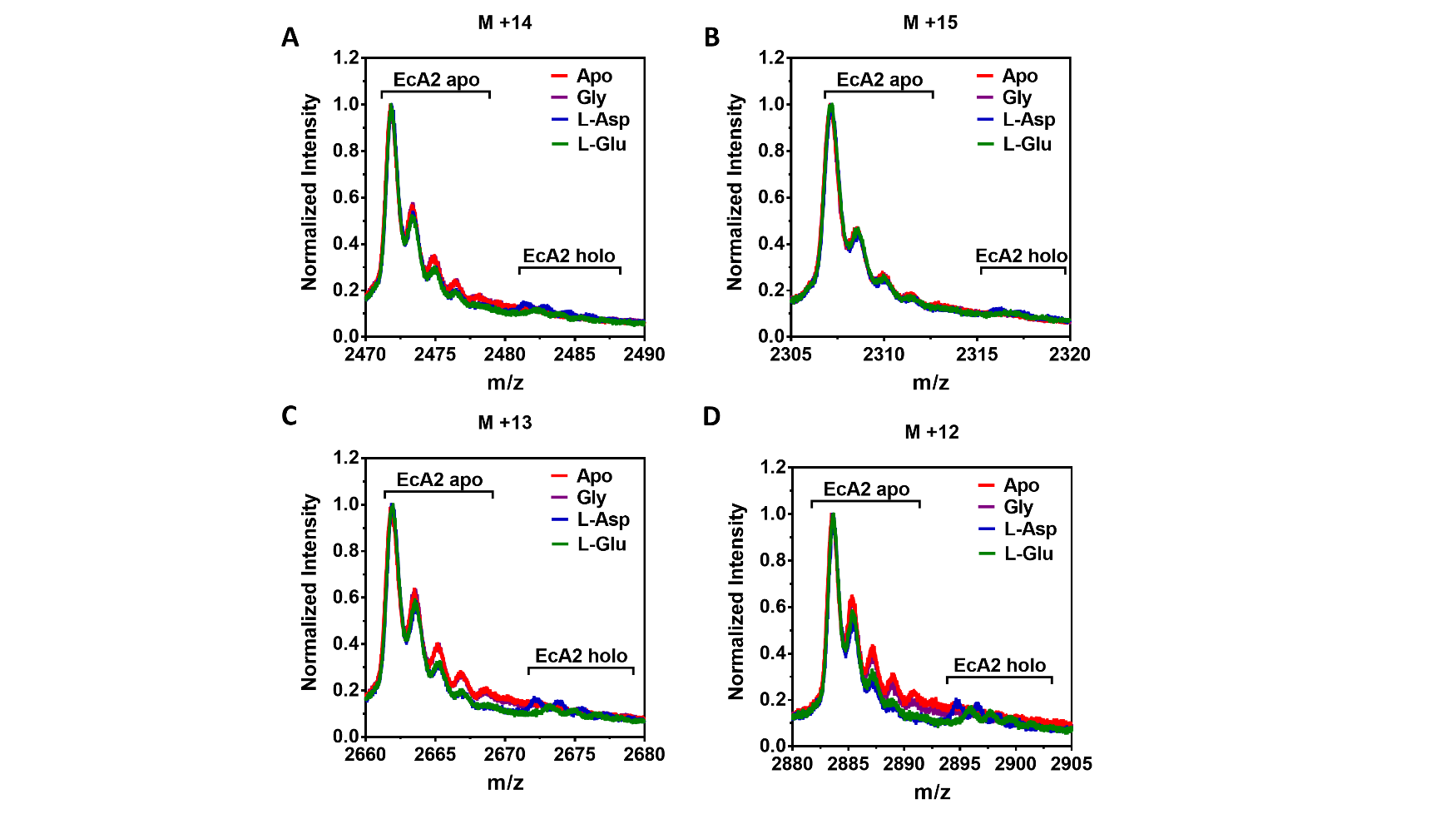

**Fig. S8.** Mapping interaction of amino acid with EcA2 by MS. The ESI-IMS-MS spectra of EcA2 were measured in the absence or presence of glycine, L-aspartate or L-glutamate, and **A.** the M+15, **B.** M+14, **C.** M+13 and **D.** M+12 charged states are shown revealing the peaks corresponding to the free and the amino acid-bound forms of the protein and salt adducts.

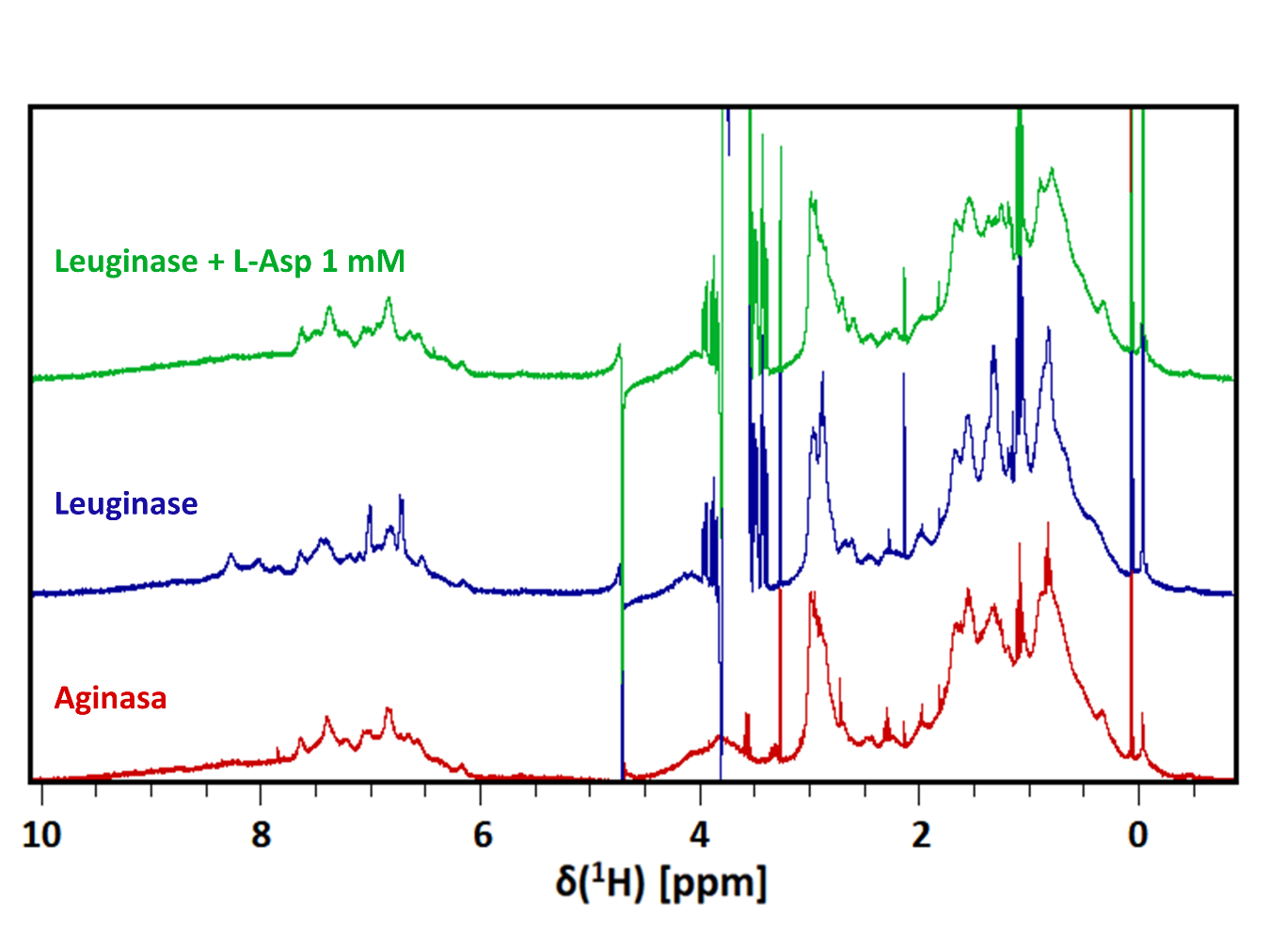

**Fig. S9.** Multiple displays of 1D ^1^H NMR spectra of Aginasa (red) and Leuginase (blue) whole formulation (320 μM) and Leuginase (320 μM) added with 1 mM of L-Asp (green). For each experiment, a total of 512 scans with 2 s of relaxation delay were collected in a 400 MHz spectrometer at 298 K. Multiple displays of NMR-titration analysis of 160 μM Leuginase supplemented with increasing concentrations of

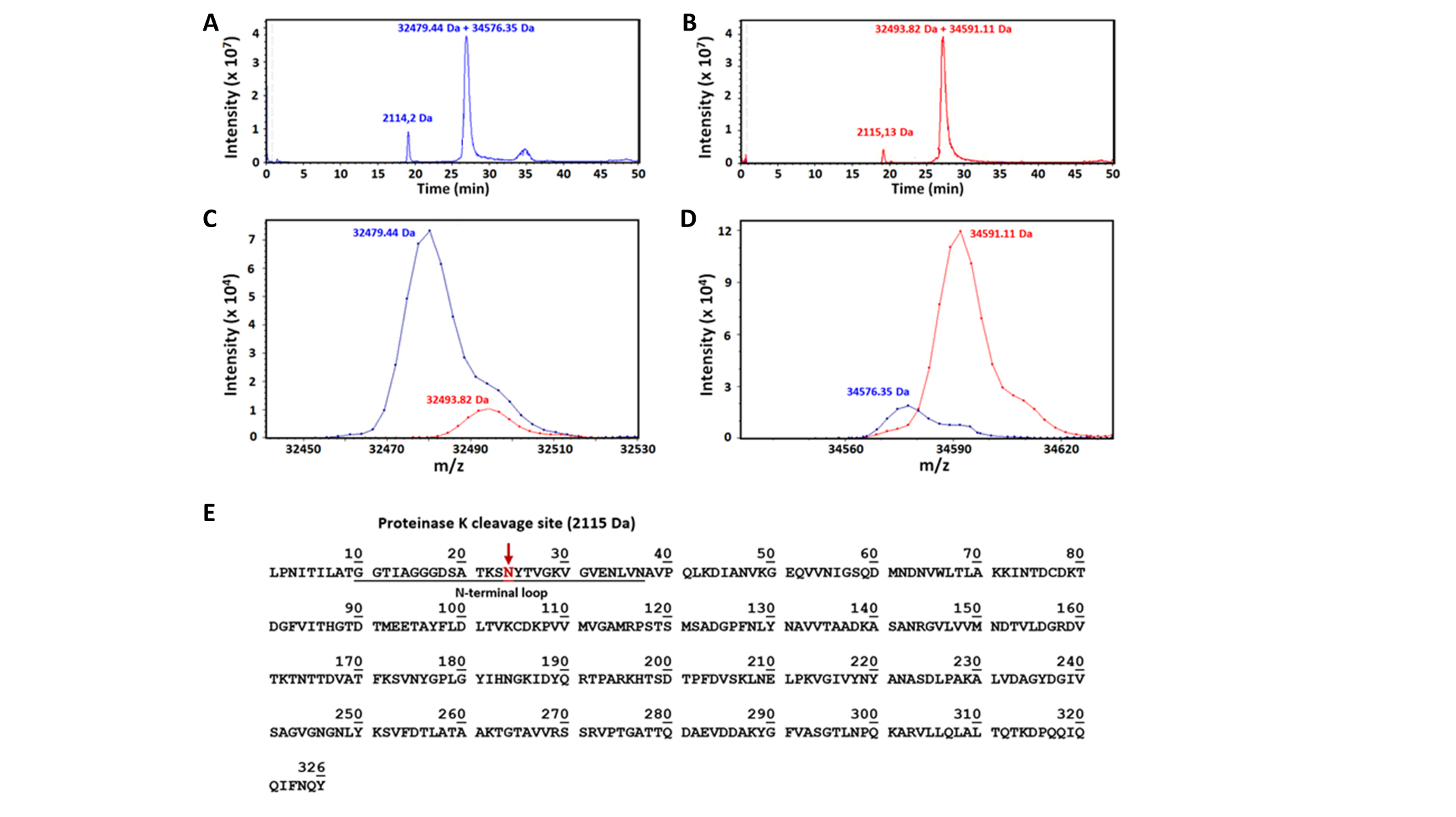

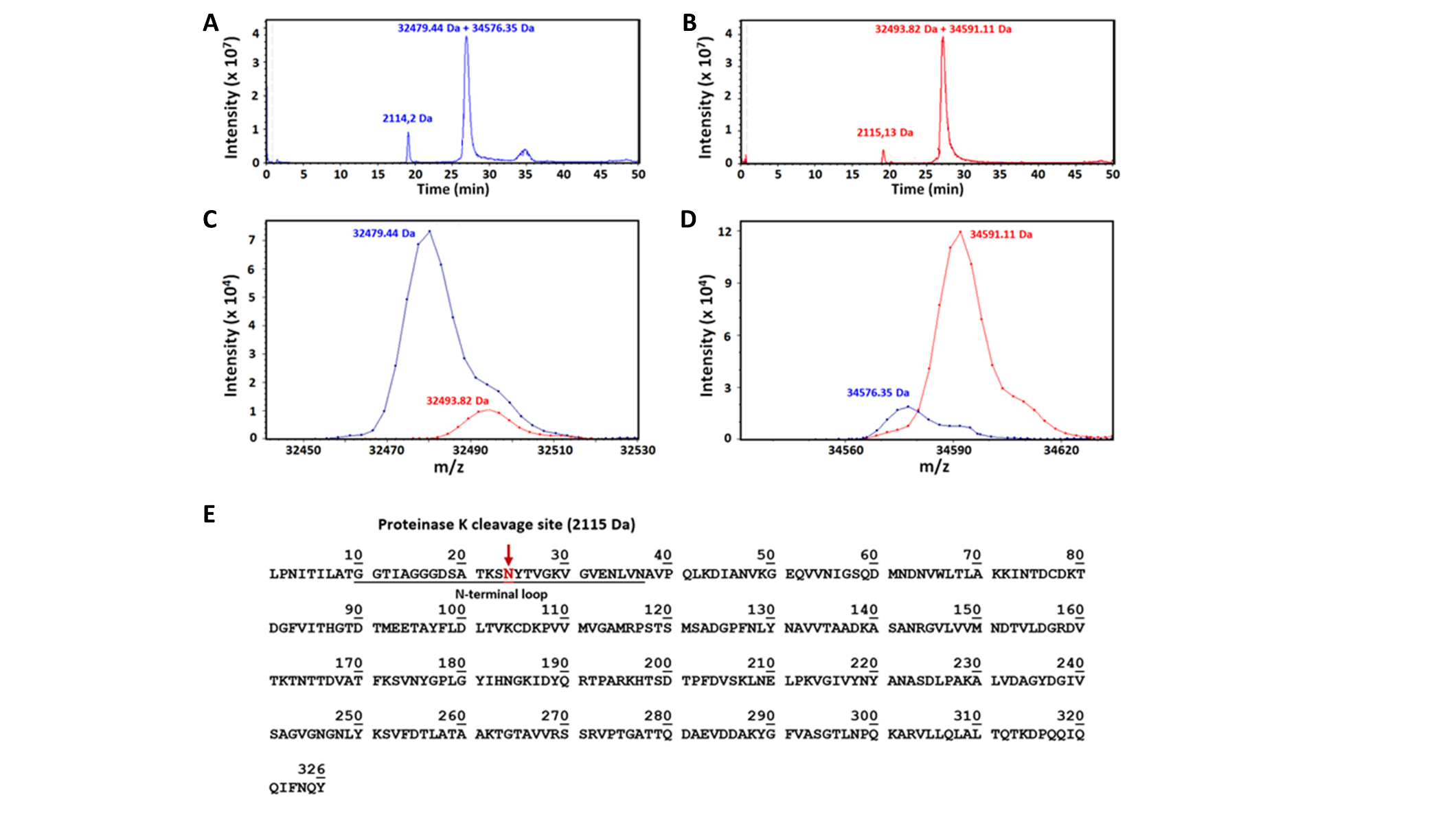

**Fig. S10.** Characterization of limited proteolysis EcA2 products by LC-MS. The total ion current (TIC) chromatograms for **A.** Leuginase and **B.** Aginasa showed a major peak corresponding to the intact proteins, along with EcA2 remaining fragment and a minor peak corresponding to the N-terminal fragment of 23 residues susceptible to proteolysis. The deconvoluted spectra revealed a mass of **C.** 34,591.11 and 32,493.82 Da for Aginasa (red) and **D.** 34,579.44 and 34,576.35 Da for Leuginase (blue), intact and proteolysis product, respectively. **E.** EcA2 amino acid sequence highlighting the suggested proteinase K cleavage site, N-terminal to N24, resulting in the 2,115 Da 23-residue fragment.

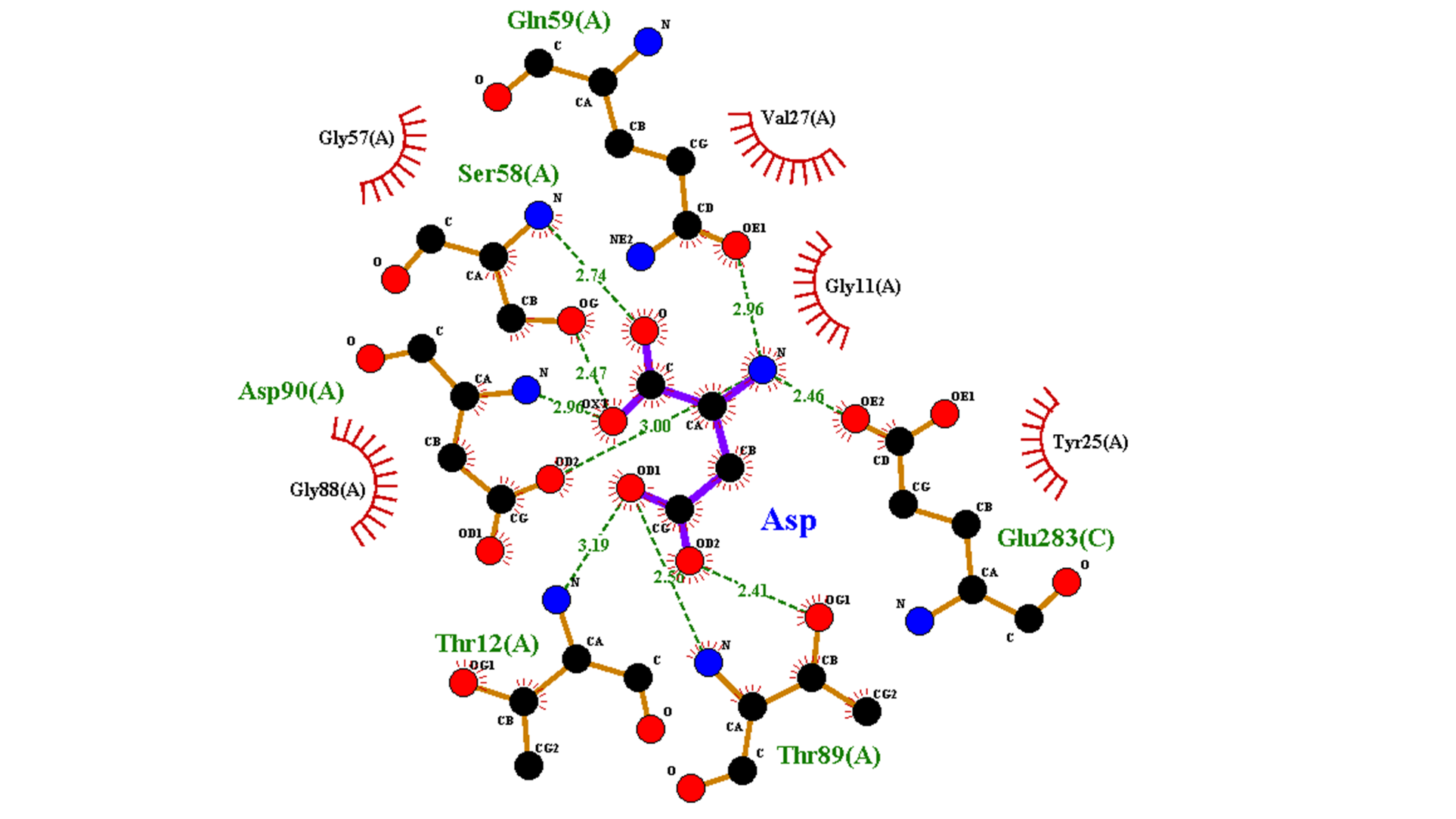

**Fig. S11.** LigPlot diagram of EcA2 catalytic site upon L-Asp binding. Ligand bonds are shown in purple and protein bonds in brown. Hydrogen bonds are represented as green dotted lines with length indicated in angstroms. Residues in hydrophobic contacts are represented by red semicircles.
